# Genetic variation in apolipoprotein A-I concentrations and risk of coronary artery disease

**DOI:** 10.1101/576504

**Authors:** Minna K. Karjalainen, Michael V. Holmes, Qin Wang, Olga Anufrieva, Mika Kähönen, Terho Lehtimäki, Aki S. Havulinna, Kati Kristiansson, Veikko Salomaa, Markus Perola, Jorma S. Viikari, Olli T. Raitakari, Marjo-Riitta Järvelin, Mika Ala-Korpela, Johannes Kettunen

**Affiliations:** Computational Medicine, Faculty of Medicine, University of Oulu, Oulu, Finland; Center for Life Course Health Research, Faculty of Medicine, University of Oulu, Oulu, Finland; Biocenter Oulu, University of Oulu, Oulu, Finland; Medical Research Council Population Health Research Unit, University of Oxford, Oxford, UK; Clinical Trial Service Unit & Epidemiological Studies Unit, Nuffield Department of Population Health, University of Oxford, Oxford, UK; National Institute for Health Research, Oxford Biomedical Research Centre, Oxford University Hospital, Oxford, UK; Medical Research Council Integrative Epidemiology Unit at the University of Bristol, Bristol, UK; Systems Epidemiology, Baker Heart and Diabetes Institute, Melbourne, VIC, Australia; Department of Clinical Physiology, Tampere University Hospital, and Finnish Cardiovascular Research Center Tampere, Faculty of Medicine and Health Technology, Tampere University, Tampere, Finland; Department of Clinical Chemistry, Fimlab Laboratories, and Finnish Cardiovascular Research Center Tampere, Faculty of Medicine and Health Technology, Tampere University, Tampere, Finland; National Institute for Health and Welfare, Helsinki, Finland; Institute for Molecular Medicine Finland (FIMM-HiLIFE), Helsinki, Finland; Diabetes and Obesity Research Program, University of Helsinki, Helsinki, Finland; Estonian Genome Center, University of Tartu, Tartu, Estonia; Department of Medicine, University of Turku, Turku, Finland; Division of Medicine, Turku University Hospital, Turku, Finland; Research Centre of Applied and Preventive Cardiovascular Medicine, University of Turku, Turku, Finland; Department of Clinical Physiology and Nuclear Medicine, Turku University Hospital, Turku, Finland; Unit of Primary Health Care, Oulu University Hospital, OYS, Oulu, Finland; Department of Epidemiology and Biostatistics, MRC-PHE Centre for Environment and Health, School of Public Health, Imperial College London, London, UK; Department of Life Sciences, College of Health and Life Sciences, Brunel University London, UK; Population Health Science, Bristol Medical School, University of Bristol, Bristol, UK; NMR Metabolomics Laboratory, School of Pharmacy, University of Eastern Finland, Kuopio, Finland; Department of Epidemiology and Preventive Medicine, School of Public Health and Preventive Medicine, Faculty of Medicine, Nursing and Health Sciences, The Alfred Hospital, Monash University, Melbourne, VIC, Australia

**Keywords:** Coronary artery disease, Apolipoprotein A-I, Mendelian randomization

## Abstract

**Rationale:** Apolipoprotein A-I (apoA-I) infusions represent a potential novel therapeutic approach for the prevention of coronary artery disease (CAD) with phase III cardiovascular outcome trials currently underway. Although circulating apoA-I levels inversely associate with risk of CAD, the evidence base of this representing a causal relationship is lacking.

**Objective:** To assess the causal role of apoA-I in CAD using human genetics.

**Methods and Results:** We identified a variant (rs12225230) in *APOA1* locus that associated with circulating apoA-I concentrations at GWAS significance (P<5×10^−8^) in 20,370 Finnish participants and meta-analyzed our data with a previous genome-wide association study of apoA-I. We obtained genetic estimates of CAD from UK Biobank and CARDIoGRAMplusC4D (totaling 122,733 CAD cases) and conducted a two-sample Mendelian randomization analysis. We compared our genetic findings to observational associations of apoA-I with risk of CAD in 918 incident CAD cases among 11,535 individuals from population-based prospective cohorts. We also summarized the available evidence from randomized controlled trials (RCTs) of apoA-I infusion therapies reporting CAD events. ApoA-I was associated with a lower risk of CAD in observational analyses (HR 0.81; 95%CI: 0.75, 0.88; per 1-SD higher apoA-I), with the association showing a dose-response relationship. Rs12225230 associated with apoA-I concentrations (per-C allele beta 0.076 SD; SE: 0.013; P=1.5×10^−9^) but not with potential confounders. In Mendelian randomization analyses, apoA-I was not related to risk of CAD (OR 1.13; 95%CI: 0.98, 1.30 per 1-SD higher apoA-I), which was different to the observational association (P-het<0.001). RCTs of apoA-I infusions did not show an effect on the risk of CAD.

**Conclusions:** Genetic evidence fails to support a cardioprotective role for apoA-I. This casts doubt on the likely benefit of apoA-I infusion therapy in the ongoing phase III cardiovascular outcome trial.

## INTRODUCTION

Large-scale cardiovascular outcome trials^1–4^ and studies of human genetics^5, 6^ do not support a causal role of high-density lipoprotein (HDL) cholesterol in coronary artery disease (CAD). However, it remains feasible that other aspects of HDL, such as the functional attributes of HDL particles, might have atheroprotective effects. The ability of HDL particles to extract cholesterol from lipid-laden cells, so-called cholesterol efflux, has recently emerged as the most prominent new measure for HDL-related atheroprotectivity.^7^ Apolipoprotein A-I (apoA-I) is a key functional apolipoprotein component of HDL particles and plays a central role in cholesterol efflux.^8^ This has led to anticipation that modification of circulating apoA-I might represent a novel therapeutic approach to the treatment and prevention of CAD. Notably, Pfizer reportedly paid over 1bn USD for the commercial rights for an apoA-I infusion technology.^9^

Recent results from phase II randomized controlled trials (RCTs) of apoA-I that used MDCO-216 (recombinant apoA-I-Milano) and CER-001 (recombinant wild-type apoA-I) infusions suggest that the initial optimism may be misplaced.^10, 11^ Trials of MDCO-216 and CER-001 failed to identify a beneficial effect of apoA-I infusion on the regression of coronary atherosclerosis as measured by intravascular ultrasonography,^10, 11^ resulting in the termination of the development of these apoA-I products. A third apoA-I infusion therapy, CSL112, representing a modified form of native apoA-I from human plasma, is currently underway in a large phase III RCT (AEGIS-II; ClinicalTrials.gov Identifier: NCT03473223), which seeks to assess the efficacy of CSL112 on the risk of cardiovascular events.^12, 13^ Prior trials have shown have shown CSL112 to be well-tolerated,^14, 15^ and capable of increasing HDL cholesterol efflux in both healthy individuals and CAD patients.^16–18^

Human genetics can yield reliable evidence on the likely efficacy of pharmacological modification of a therapeutic target on risk of disease. For example, genetic variants in *HMGCR* and *PCSK9* reliably elucidate the effects of statins and PCSK9 inhibition on circulating metabolic markers (including lipoproteins and their accompanying lipids) and risk of CAD.^19–21^ In this study, we used human genetics in five population-based cohorts with detailed lipoprotein profiling together with large-scale genome-wide data to identify a robust genetic instrument for apoA-I in order to provide reliable estimates of the potential causal effects of apoA-I on risk of CAD. We also compared our genetic results to observational associations of apoA-I and the risk of incident CAD in population-based cohorts and analyzed data from previously published RCTs of apoA-I infusion therapies reporting CAD events.

## METHODS

### Datasets

We used individual participant data including genotyping and lipoprotein profiling from five Finnish population-based cohorts, totaling up to 20,370 individuals (Supplemental Table 1): Northern Finland Birth Cohorts^22, 23^ 1966 (NFBC66) and 1986 (NFBC86), the National FINRISK^24^ studies 1997 (FINRISK97) and 2007 (FINRISK07) and the Cardiovascular Risk in Young Finns Study^25^ (YFS). Details of the studies are provided in the Supplement. The genetic studies were conducted using data from these five Finnish cohorts. Observational estimates were assessed in the FINRISK cohorts (Supplemental Table 2). In addition, we used summary-level data from large consortia^26–29^ as described in the Supplement.

### Identifying genetic instruments for the circulating apoA-I concentration

To find genetic variants that mimic the effect of apoA-I infusion therapies, we focused on *cis*-acting protein quantitative trait loci (pQTLs) in the *APOA1* locus, which contains the gene encoding apoA-I. We concentrated on the 1-Mb region flanking the *APOA1* gene to find genetic variants associated with circulating apoA-I concentrations. Our initial approach was to count the number of single-nucleotide polymorphisms (SNPs) in this region and use this value to correct the multiple testing through a conventional Bonferroni approach, which corresponded to 7,210 SNPs and a Bonferroni-adjusted P-value of <7×10^−6^. However, the top SNP superseded this threshold and associated with apoA-I at conventional levels of GWAS significance (P<5×10^−8^). To identify whether more than one cis-pQTL associated with apoA-I, we repeated the analysis of apoA-I concentrations on SNPs in/around *APOA1* after conditioning on our top variant. We compared our results to those of a previous GWAS of apoA-I^30^ and prioritized variant rs12225230 as the genetic instrument for apoA-I; this SNP was reported to be associated with apoA-I in the previous GWAS^30^ and summary statistics were available from this study. No additional conditionally-independent cis-pQTLs were identified. Details of the analyses and selection criteria are described in the Supplement.

### Assessment of pleiotropy of the genetic instrument

To comprehensively annotate the metabolic effects of the genetic instrument for apoA-I, we investigated the effects of the genetic variant across a wide range of lipoprotein lipids and metabolites analyzed by nuclear magnetic resonance (NMR) spectroscopy (228 measures, Supplemental Table 4).^31, 32^ The metabolites were first inverse-rank normalized, then adjusted for age, sex, and ten first principal components, and inverse rank-based normal transformation was used to transform the resulting residuals to a normal distribution to follow the normalization procedure of our previous GWAS.^33^ For the association of rs12225230 with metabolite levels, we considered P values <0.002 as an evidence of association (Bonferroni correction with 22 principal components that explained >95% of total variation in the levels of the metabolic measures). To facilitate comparison across multiple continuous measures, effect sizes are reported as standard deviation (SD) units per allele and illustrated as forest plots drawn with R, v. 3.4.3. To identify potential pleiotropic effects of the variant, we also conducted a phenome-wide association analysis in PhenoScanner.^34^

### Observational association of apoA-I with risk of incident coronary artery disease

We estimated the association of serum apoA-I levels with risk of incident CAD (defined as MI, coronary revascularization, or death from CAD; see Supplement for details) in FINRISK97 (N=7,133; number of incident CAD events 743 during 18-year follow-up) and FINRISK07 (N=4,402; number of incident CAD events 175 during 8-year follow-up).^24^ To place the apoA-I association with CAD into context of other lipid measures, we also estimated the prospective associations of apolipoprotein B (apoB), HDL cholesterol and LDL cholesterol with CAD in the FINRISK97 cohort. Effect sizes were estimated using Cox proportional-hazard model and are reported in HR per SD difference in the circulating concentration. Two models were used: a minimally adjusted model with age and sex as covariates, and a fully adjusted model including a comprehensive set of covariates (age, sex, LDL cholesterol, body-mass index, systolic blood pressure, type 2 diabetes, smoking and alcohol consumption). The cohorts were meta-analyzed using inverse-variance fixed-effect modeling. All analyses were conducted using R, v. 3.4.3.

### Analysis of randomized clinical trials of apoA-I infusion therapies

To quantify the effect of apoA-I infusion from recent RCTs, we first identified trials in PubMed using the following search criterion: “apolipoprotein A-I infusion randomized clinical trial” (on 10/01/2018). We further assessed the articles identified in PubMed to identify placebo-controlled RCTs that reported major adverse cardiovascular events (MACEs) as endpoints (Supplemental Table 5). This identified the CHI-SQUARE^35^ trial (3, 6, or 12 mg/kg infusions of CER-001 weekly for six weeks) and the AEGIS-I^15^ trial (2 or 6 g infusions of CSL112 weekly for four weeks). The analyzed end point was MACE defined as cardiac arrest, death, non-fatal MI, non-fatal stroke, coronary revascularization, or hospitalization for unstable angina or heart failure in the CHI-SQUARE trial; and as cardiovascular death, nonfatal MI, ischemic stroke or hospitalization for unstable angina in the AEGIS-I trial. The total number of MACE cases was 55 in CHI-SQUARE (total number of trial participants=486) and 74 in AEGIS-I (total N=1,258). We extracted estimates corresponding to all doses of apoA-I infusion therapies and calculated relative risks for MACEs for each dose separately.

### Assessing causality

We assessed causality between the exposure (circulating apoA-I levels) and outcome (CAD) utilizing the Mendelian randomization (MR) framework (Supplemental Figure 1).^36,37^ We used a two-sample MR approach where summary data of the exposure (i.e., SNP to apoA-I level) and outcome (i.e., SNP to CAD) originate from different sources.^36,37^ Rs12225230 located in the *APOA1* locus was used as the genetic instrument. This approach included the use of the following estimates: [1] per-allele effect estimate (in g/L) for the association between rs12225230 and apoA-I (meta-analyzed result between the Finnish individual participant data (IPD) and a prior GWAS^30^; total N=37,093); [2] per-allele effect estimates with risk of CAD from a large GWAS, i.e., meta-analysis of the UK Biobank with CARDIoGRAMplusC4D^29^ (case N=122,733). To obtain a causal estimate and the corresponding standard error for the association between apoA-I and CAD, we used the formula described by Burgess *et al^38^* that collapses to Wald ratio for a single variant. To facilitate comparison with observational estimates, the causal estimate of apoA-I on risk of CAD was scaled to SD units, using SD of 0.225 g/L (the SD of apoA-I in our largest cohort, FINRISK97) and assessed evidence of heterogeneity between the causal and observational estimates using Cochrane’s Q statistic. In sensitivity analyses, we repeated the MR analysis using the per-allele effect estimate between rs12225230 and apoA-I using values derived from just the Finnish population cohorts and just the prior GWAS.^30^ We report our power to detect various effect estimates of apoA-I and risk of CAD in the MR analyses in Supplemental Table 3.

## RESULTS

### Identification of genetic variants associated with circulating apoA-I concentrations

Multiple SNPs in/around the *APOA1* locus were associated with apoA-I serum concentration (Figure 1, Supplemental Table 6) with the association peak spanning a 466-kb region. We identified the missense variant rs12225230 to associate with apoA-I concentrations at GWAS significance (per-allele beta 0.076 SD, P=1.5×10^−9^), which was in linkage disequilibrium (LD) with the top SNP rs625145 (*D*’=1.00, *r*^2^=0.91). This missense variant was previously shown to be associated with apoA-I serum concentration^30^ and is mapped to the *SIK3* gene encoding SIK family kinase 3, located 20 kb upstream of *APOA1*. When we conditioned on rs12225230, there were no additional associations (Supplemental Figure 2), indicating that rs12225230 or SNPs in LD with this SNP are likely the principal drivers of the apoA-I association in this locus. We estimated the effect of the minor allele of rs12225230 on apoA-I levels to be 0.0158 g/L (SE 0.0030). In meta-analysis of our cohorts with a previous GWAS of apoA-I (reporting 0.0320 g/L [SE 0.0033] increment per minor allele of rs12225230)^30^ the effect of the minor allele C of rs12225230 was estimated to be 0.0230 g/L (SE 0.0022; P=1.5×10^−25^), corresponding to an F-statistic of 109.

**Figure 1.**
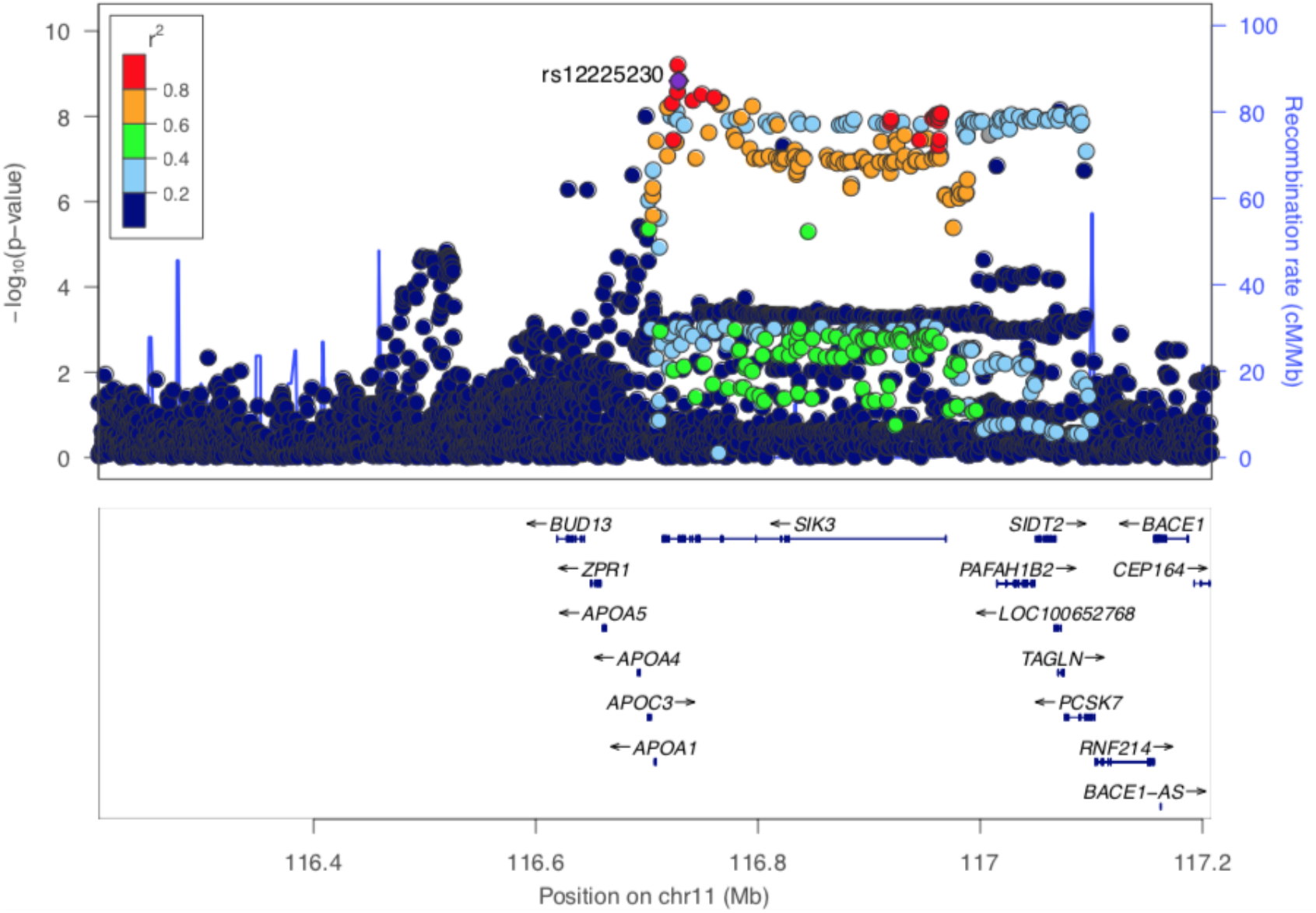
Association of SNPs in the *APOA1* locus with circulating concentrations of apoA-I. Each dot represents the association of a single genetic variant with apoA-I levels in metaanalysis of five Finnish population cohorts (−log10 of P value shown on the y axis and chromosomal position on the x axis). The 500-kb region flanking *APOA1* is shown. Rs12225230, highlighted in violet and was used as an instrument in this study, is robustly associated with apoA-I (P=2×10^−9^) and is in LD with the top variant rs625145 (P=6×10^−10^).

When we further annotated this SNP, rs12225230 mapped to a region containing regulatory chromatin states and enhancer histone marks in several tissues, including the liver. In large eQTL data, there were no significant eQTLs in the *APOA1* locus in the liver or small intestine.^26,39^ According to mQTL data^40^, rs12225230 is associated with the methylation status of nearby CpG methylation sites. Furthermore, rs12225230 is in LD with rs625145 (top SNP in our apoA-I GWAS), and methylation status of CpG sites associated with this SNP was previously reported to be causally associated with apoA-I concentrations.^41^

We assessed whether rs12225230 represents an unbiased instrument to mimic the effect of apoA-I increasing therapies by characterizing the effects of rs12225230 on metabolic measures (Figure 2, Supplemental Figure 3). We detected that, in addition to associating with higher apoA-I, the minor allele of rs12225230 was associated with higher levels of multiple HDL-related measures, including, e.g., concentration of large, medium and small HDL particles (L-HDL-P, beta 0.055 SD, P=1.2E-05; M-HDL-P, beta 0.061 SD, P=1.2E-06; S-HDL-P, beta 0.055 SD, P=1.2E-05), and HDL cholesterol (beta 0.070 SD, P=2.5E-08). In addition, rs12225230 was associated, e.g., with fatty acid measurements (e.g., polyunsaturated fatty acids, beta 0.048 SD, P=1.2E-04); associations with the fatty acids likely reflect their associations with multiple HDL-related traits. In contrast, rs12225230 was not associated with the majority of apoB related traits (Supplemental Figure 3).

**Figure 2.**
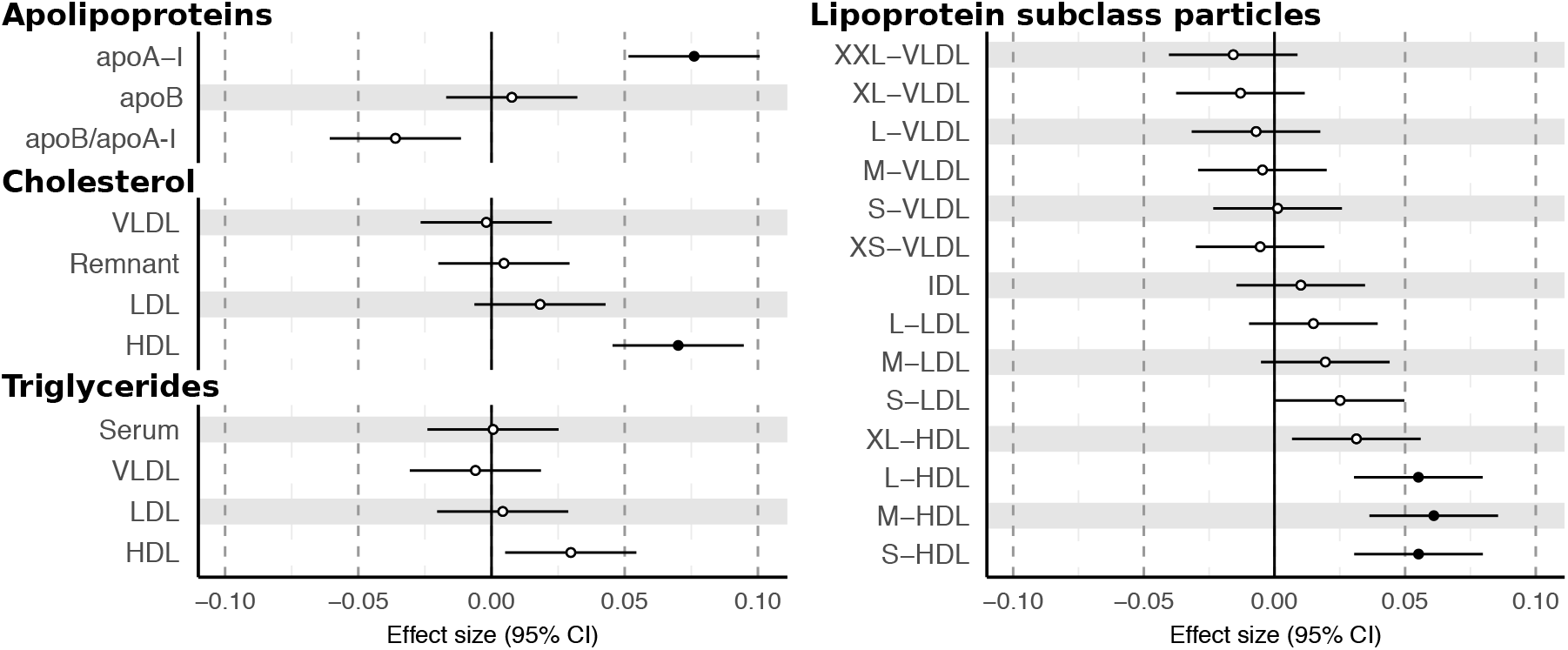
Association of rs12225230 with key lipoprotein-related concentration measures. Remnant cholesterol refers to cholesterol carried in VLDL and IDL particles. Effect sizes are estimated as SD differences in metabolite concentrations per rs12225230-C allele. Closed symbols, P<0.002 (significant association); open symbols P≥0.002. Abbreviations: apoA-I, apolipoprotein A-I; apoB, apolipoprotein B; VLDL, very low-density lipoprotein, LDL, low-density lipoprotein; HDL, high-density lipoprotein; IDL, intermediate-density lipoprotein.

We further investigated whether the association of rs12225230 with apoA-I level could be confounded by associations with other phenotypes. To this end, we screened publicly available genotype-phenotype associations using PhenoScanner.^34^ The associations of rs12225230 were not likely confounded by conventional risk factors (see Supplemental Methods, Supplemental Table 7, Supplemental Figure 4).^42, 43^

### Observational associations of apoA-I levels with incident coronary artery disease

We estimated the prospective observational associations of apoA-I with risk of CAD in the FINRISK97 cohort (N=7,133, with 743 incident CAD events), and compared the association of apoA-I to the corresponding CAD associations for apoB, HDL and LDL cholesterol. ApoA-I (HR 0.86 per 1-SD higher apoA-I; 95%CI 0.79-0.93, Figure 3) had a similar magnitude of association with CAD as did HDL cholesterol (HR 0.79 per 1-SD higher HDL-C; 95%CI 0.72-0.86), apoB (HR 0.84 per 1-SD lower apoB; 95%CI 0.79-0.91) and LDL cholesterol (HR 0.88 per 1-SD lower LDL-C; 95%CI 0.82-0.95). The prospective association of apoA-I with risk of CAD was similar in minimally (age and sex as covariates) and fully (age, sex, body mass index, systolic blood pressure, type 2 diabetes, smoking and alcohol consumption as covariates) adjusted models (Supplemental Table 8). ApoA-I showed a dose-response relationship, with the risk of CAD being HR 0.57 (95%CI 0.45-0.75) comparing the highest to lowest quintiles of apoA-I (Figure 3B).

**Figure 3.**
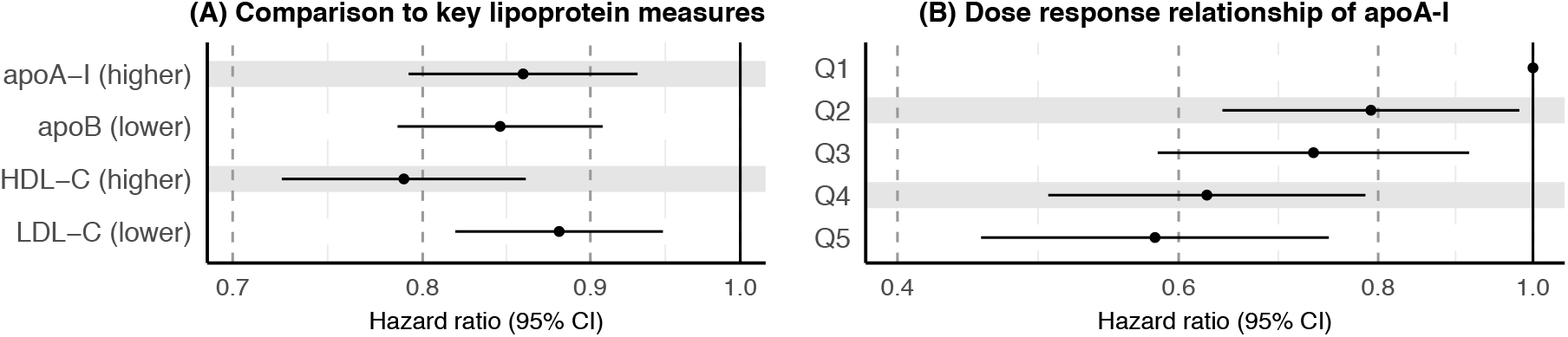
Observational associations of apoA-I and risk of coronary artery disease. **Panel A:** the association of apoA-I with CAD risk was compared to those of key lipoprotein measures (apoB, HDL and LDL cholesterol). Note, the effects of apoB and LDL cholesterol are inversed; i. e., the effects are shown per 1-SD reduction in apoB and LDL cholesterol. Associations were assessed with 743 cases and 6,390 controls in FINRISK97, with adjustment for age, sex, body-mass index, systolic blood pressure, type 2 diabetes, smoking and alcohol consumption. Associations are presented as HR (95%CI) per 1-SD lipoprotein measure. **Panel B:** Dose-response association of apoA-I with risk of CAD. Associations were additionally adjusted for LDL cholesterol. Effect estimates are presented per quintile of apoA-I with the lowest quintile (Q1) used as the reference category.

In meta-analysis of the FINRISK97 and FINRISK07 cohorts (total 918 incident CAD cases), higher concentrations of apoA-I associated with a lower risk of CAD (HR 0.81 per 1-SD higher apoA-I, 95%CI 0.75-0.88) after adjustment for age, sex, LDL cholesterol, body-mass index, systolic blood pressure, type 2 diabetes, smoking and alcohol consumption. The associations were similar in minimally and fully adjusted models and in both cohorts (Supplemental Figure 5). ApoA-I showed only low correlation (r between −0.3 and 0.3) with the potential confounders included as covariates in the models (Supplemental Figure 6).

### Effects of apoA-I infusion therapies on major adverse cardiovascular events

We appraised the evidence base on the relationship of apoA-I infusion therapies and risk of cardiovascular disease in two phase II RCTs reporting major adverse cardiovascular event (MACE) endpoints: the CHI-SQUARE^35^ and AEGIS-I^15^ trials. None of the apoA-I infusion doses reduced the risk of MACEs: while dose-specific effect estimates had wide 95% confidence intervals, it is noteworthy that at all doses investigated, the point estimates were on the opposite side of the null that would indicate a potential protective effect (Supplemental Table 9).

### Evaluation of the causal role of serum apoA-I in coronary artery disease

Using data from a large GWAS of coronary artery disease including 122,733 CAD cases in 547,261 individuals (meta-analysis of UK Biobank CAD GWAS with CARDIoGRAMplusC4D^29^), rs1222530 was not associated with risk of CAD (per-allele log-odds 0.0124, SE 0.0073, P=0.090). Similarly, no significant association of rs1222530 was reported in any of the large GWAS for cardiovascular phenotypes, including CAD, MI or stroke (Supplemental Table 10). Using the rs1222530-apoA-I and rs1222530-CAD associations, we did not find evidence to support a causal role of apoA-I with risk of CAD: the causal estimate (calculated based on meta-analysis of our own GWAS and a previous GWAS of apoA-I^30^) for CAD was an OR of 1.13 per 1-SD higher apoA-I level, 95%CI 0.98-1.30 (Figure 4, Supplemental Figure 7) which differed (P-heterogeneity<0.001) to the corresponding fully-adjusted observational estimate (HR 0.81; 95%CI: 0.75-0.88). Causal estimates for CAD were similar when calculated based on the apoA-I effect estimates from the previous GWAS^30^ (OR 1.09, 95%CI 0.99-1.21) and our own GWAS (OR 1.19, 95%CI 0.97-1.46).

**Figure 4.**
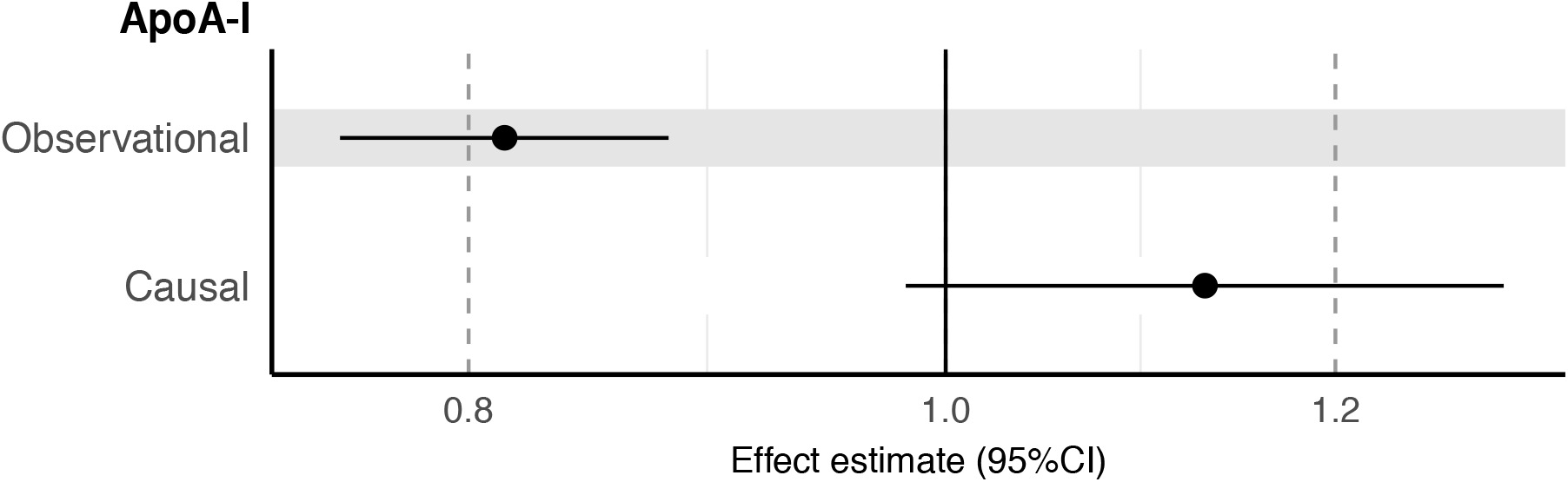
Observational and causal effects of circulating apoA-I levels and risk of coronary artery disease. Observational estimate was assessed in the FINRISK97 and FINRISK07 cohorts (918 coronary artery disease cases among 15,353 individuals) with age, sex, LDL cholesterol, body-mass index, systolic blood pressure, type 2 diabetes, smoking and alcohol consumption as covariates. Causal estimate was calculated based on association of rs12225230 with apoA-I levels (effect estimate from meta-analysis of five Finnish cohorts and a previous GWAS^30^, total N=37,493) and coronary artery disease (effect estimate from meta-analysis of the UK Biobank and CARDIoGRAMplusC4D^29^; 122,733 CAD cases among 547,261 individuals). Effect estimates correspond to the risk (observational association: hazard ratio, HR; causal estimate: odds ratio, OR) of CAD per 1-SD higher apoA-I (equivalent to 0.225 g/L apoA-I).

## DISCUSSION

Our findings do not provide genetic support for the hypothesis that circulating apoA-I concentrations play a protective role in the etiology of CAD. Together with previous genetic and randomized controlled trial evidence indicating that HDL cholesterol does not have a protective effect on risk of CAD,^1,2,5,6^ these results suggest that apoA-I is unlikely to represent a valid therapeutic approach for the prevention and treatment of CAD. The implication of these findings is that the ongoing AEGIS-II phase III cardiovascular outcome trial of CSL112 is unlikely to show evidence of vascular benefit.

HDL cholesterol and apoA-I are among the serum lipoprotein measures showing strongest inverse associations with cardiovascular disease in epidemiological studies.^44, 45^ However, large phase III cardiovascular outcome RCTs using HDL cholesterol increasing therapies have failed to identify that the risk of CAD is proportionate to the amount by which HDL cholesterol is increased,^1, 2^ and, consistently, studies of human genetics find that the association of HDL cholesterol with CAD is not causal.^5, 6^ In the REVEAL trial of the cholesteryl ester transfer protein (CETP) inhibitor, anacetrapib, although treatment with anacetrapib did increase HDL cholesterol, the cardiovascular benefit was proportionate to the degree of apoB lowering, rather than to the effect on HDL cholesterol.^3^ Here, we add further evidence showing that targeting HDL-related pathways through increases in apoA-I is unlikely to be effective in lowering risk of CAD. We show that long-term genetically increased circulating apoA-I levels do not protect from CAD, a finding consistent with the cardiovascular associations in two trials of apoA-I infusion products (MDCO-216 and CER-001) that led to the termination of development of those products.^10, 11^ Thus, randomized evidence of both the short-term, transient increase in circulating concentration of apoA-I arising from apoA-I infusions and the life-long increase caused by genetic variants fails to provide evidence in support of a protective role of apoA-I in CAD.

This study has several strengths. The SNP used to mimic the effect of apoA-I-increasing therapy, rs12225230, was identified in a locus-specific analysis of SNPs in/around *APOA1* in meta-analysis of five Finnish cohorts in our analyses and the same variant was identified in a prior GWAS of apoA-I^30^ with the mechanism thought to be related to alterations in methylation of regulatory sites. Having an F-statistic of 100, and identified through scanning genetic variants in the cis-encoding locus, the SNP represents a robust and valid genetic instrument for apoA-I. By only considering *cis*-acting pQTL SNPs from the *APOA1* locus, we minimized the potential for confounding by *trans*-acting variants, since the use of *cis*-acting pQTLs as instrumental variables for proteins represents among the most reliable of approaches to Mendelian randomization.^37^ In addition, we carefully assessed the effects of rs12225230 on a detailed panel of serum metabolic measures as a means of providing comprehensive evidence against potential confounding by circulating metabolic markers other than those related to HDL pathways, and supplemented this by examining potential associations using publically available genetic repositories. For all analyses, we utilized large, well-characterized Finnish population cohorts and the largest available international GWAS consortia data for CAD to maximize statistical power. Taken together, our statistical and genetic approach yielded reliable estimates of the causal effect of apoA-I on risk of CAD and facilitated the discovery that, unlike the observational association, which is vulnerable to confounding and bias, apoA-I is unlikely to play a causal role in CAD.

The main focus of investigation was to provide genetic evidence on the hypothesis of whether apoA-I infusions protect from CAD, and to do so, we developed a genetic instrument that strongly and specifically associates with circulating apoA-I concentrations as the primary exposure. However, the rationale behind apoA-I-infusion therapies is that increases in apoA-I ought to promote cholesterol efflux from arterial wall macrophages, a process that can be quantitatively measured by HDL cholesterol efflux capacity (HDL-CEC). Similar to HDL cholesterol and apoA-I, HDL-CEC is inversely associated with risk of CAD.^7, 46^ Whether or not HDL-CEC is modified by apoA-I in a cause and effect manner is not currently known. The AEGIS trial showed that apoA-I infusion therapy with CSL112 led to higher concentrations of apoA-I and higher HDL-CEC,^15, 18^ which suggests that higher HDL-CEC may represent a target-mediated effect of elevations in apoA-I. If this were true, then we would expect that our genetic instrument for higher apoA-I would also lead to higher HDL-CEC. This therefore means that we would expect that any effect of apoA-I infusion therapy on HDL-CEC ought to be represented by our genetic instrument.

A common criticism of drug-target Mendelian randomization analyses is that the genetic association with the biomarker of interest (in this case apoA-I) is typically small, and thus meaningful deductions cannot be made about the likely impact of modifying apoA-I through a therapeutic (such as an infusion), where the effect of the intervention on the biomarker of interest is typically much larger. The counterargument is that atherosclerosis is a life-long disease, and exposure to a small but consistent genetic elevation in apoA-I over several decades ought to lead to a lower risk of vascular disease if apoA-I is truly cardioprotective. It is this ‘cumulative’ exposure in the setting of a disease with a long latency period (such as cardiovascular disease) which means that ‘physiological’ risk factors such as LDL cholesterol tend to have much larger effects on risk of CVD from Mendelian randomization than from conventional observational studies. ^37^ In our study, we had very good power (~80%) to detect an effect estimate from Mendelian randomization which was similar to that which we identified from our observational analyses. If apoA-I were truly causal, we would expect the effect estimate representing the actual causal estimate to be of greater magnitude (e.g. odds ratios of 0.7 or lower) for which we had excellent power. Our genetic analyses might give rise to a false negative in the context of 2-sample MR using non-overlapping datasets if we had a weak instrument, however our F-statistic was over 100, making this unlikely. A false negative might also arise were there to be a physiological threshold above which apoA-I is cardioprotective but below which it has no effect. To investigate the plausibility of such a threshold effect, our observational analyses, in which we generated quintiles based on measured values of apoA-I, showed a clear dose-response relationship between apoA-I and risk of vascular disease, which argues against such a threshold effect.

This study provides an example of using human genetics to guide the development of therapeutic targets, and predict the effects of phase III cardiovascular outcome trials. We note the similarity to other ‘negative’ MR studies of secretory phospholipase IIA-A_2_^47^ and Lp-PLA_2_^48^, both of which were the focus of large-scale phase III cardiovascular outcome trials. In both cases, reliable genetic evidence might have prevented vast sums of money from being spent on phase III cardiovascular outcome trials of therapies modifying drug targets that were destined to fail due to lack of target-mediated efficacy. While the biological plausibility of the hypothesis that apoA-I infusion might lower risk of CAD through increased reverse cholesterol transport is biologically appealing, our genetic evidence indicates that the ongoing AEGIS-II phase III trial is unlikely to succeed in lowering the risk of CAD. The cumulative evidence therefore brings into question whether pharmaceutical research and development should continue to invest in developing apoA-I infusion therapies for the purposes of preventing CAD.

In conclusion, our findings do not support the hypothesis that increasing circulating apoA-I concentrations represents a valid approach for the prevention of CAD. These results add to the burgeoning evidence that refutes a protective role of HDL and HDL-related phenotypes in the etiology of CAD.

## NON-STANDARD ABBREVIATIONS AND ACRONYMS

apoA-I: , apolipoprotein A-I;
CAD: , coronary artery disease;
eQTL: , expression quantitative trait locus;
MACE: , major adverse cardiovascular event;
MR: , Mendelian randomization;
NMR: , nuclear magnetic resonance;
pQTL: , protein quantitative trait locus;
SNP: , single-nucleotide polymorphism.

