## Supplementary material for "Genetic variation in apolipoprotein A-I concentrations and risk of coronary artery disease": Online Supplement

### SUPPLEMENTAL MATERIAL

### Detailed Methods

#### Description of the Finnish population cohorts

***Ethical approvals.*** The FINRISK, NFBC and YFS studies were approved by the following ethical committees: the National Public Health Institute, Helsinki, Finland, and Coordinating Ethical Committee of the Helsinki and Uusimaa Hospital district; Northern Ostrobothnia Hospital District, Finland; and the five universities with medical schools in Finland. Written informed consent was obtained from all participants.

***FINRISK cohorts.*** The study protocol for the population-based FINRISK studies has been described before.[1] In summary, FINRISK surveys have been conducted every five years since 1972 to study the risk of chronic diseases. A random sample of 24-74-year-old inhabitants was selected from different regions in Finland for each survey that included a questionnaire and a clinical examination (including blood sample collection); the data were linked to national registers of health outcomes. The present study included eligible individuals from the 1997 (N=6,643 in genetic analyses; N=7,133 in observational analyses) and 2007 (N=3,896 in genetic analyses; N=4,402 in observational analyses) surveys (Supplemental Tables 1 and 2). FINRISK97 serum samples were taken during the clinical visit after >4-hour fasting. FINRISK07 samples were taken from the 8-hour fasting subset of DIetary, Lifestyle, and Genetic determinants of Obesity and Metabolic syndrome (DILGOM) part of the 2007 survey[2] Genotyping of the FINRISK samples was conducted in various batches and arrays: PredictCVD (genotyped by HumanOmniExpress-12v1H), SANGER1 (genotyped by HumanCoreExome-12-v1-0-D), PSYCHO (genotyped by PsychArrayB), MRPRED (genotyped by HumanCoreExome-24v1-0A), SUMMIT (HumanOmniExpress-12v1H), MIGEN (genotyped by Affymetrix 6.0) and COROGENE (genotyped by Human 610- Quadv1H). Imputation (against the 1000genomes phase 1v3 reference genome using IMPUTE2) of the batches was performed separately, and the batches were later combined for analysis using batch as a covariate.

***Northern Finland Birth Cohorts (NFBC).*** NFBC studies are longitudinal birth cohorts that were established to study factors related to preterm birth and consequent morbidity in the northern part of Finland (provinces of Oulu and Lapland). NFBC66 covers 96% of all eligible live births during 1966 in this region (altogether 12,231 children; 12,058 alive).[3] The present study included eligible individuals from NFBC66 at the age of 31 years (N=4,668; Supplemental Table 1). NFBC86 covers 99% of all deliveries in this region during July 1985-June 1986 (altogether 9,423 births and 9,479 children).[4] Mothers and children of both cohorts have been followed-up since enrollment of the mothers at the first antenatal clinic visit (10-16$^{th}$ week). Clinical examination and serum sampling were conducted for NFBC86 participants (N=6,621) at the age of 15-16 years in 2001-2002; attendees in the 16-year field study (71% of invited participants) were representative of the original cohort. The present study included eligible individuals from NFBC86 (N=3,215). Genotyping of the NFBC66 and NFBC86 samples was performed with the following arrays: Illumina Infinium 370cnvDuo and Illumina HumanOmniExpressExome. Imputation was performed following the HRC imputation pipeline. Fasting serum samples were used for NMR metabolomics.

***Cardiovascular Risk in Young Finns Study (YFS).*** The study protocol for the prospective, population-based YFS cohort study has been described before.[5] In summary, YFS study was performed at five medical schools (Helsinki, Kuopio, Tampere, Turku and Oulu) in Finland to study levels of cardiovascular risk factors in children and adolescents in different regions of the country. The fasting serum samples used in this study were collected in 2007 follow-up and altogether 1,948 YFS participants were included (Supplemental Table 1). Genotyping of the YFS samples was performed with Illumina670K array, and imputation followed the HRC imputation pipeline.

**Diagnostic criteria for coronary artery disease cases in the FINRISK97 and FINRISK07 cohorts**

Coronary artery disease (CAD) was defined as myocardial infarction (MI), a coronary revascularization procedure, or death from coronary heart disease, i.e., falling into any of the following categories:

- I21 or I22 (**ICD-10**) / 410 (**ICD-8/9**) as the direct or as a contributing cause of death or I20-I25 (**ICD-10**) /410-414 (**ICD-9**) as the underlying cause of death
- I21 or I22 (**ICD-10**) / 410 (**ICD-8/9**) as the main or secondary diagnosis at hospital discharge.
- Coronary bypass surgery or coronary angioplasty at hospital discharge or identified from the Finnish registry of invasive cardiac procedures.

**Selection of genetic variants for apoA-I instrument and functional annotation**

Although several loci have been associated with apoA-I level in prior GWASs (*ABCA1, APOA1, APOE, GCKR, CETP, HNF4A, LIPC, LIPG, LPL*),[6-11] most of these are trans-pQTL and therefore may be non-specific for apoA-I thus rendering them potentially unsuitable for inclusion in the genetic instrument due to genetic pleiotropy.[12] Therefore, we concentrated on the *APOA1* locus. ApoA-I concentrations were first inverse-rank normalised, then adjusted for age, sex, and the first ten principal components, and inverse rank-based normal transformation was used to transform the resulting residuals to a normal distribution to follow the normalization procedure of our previous GWAS.[6] Association analysis with the normalized residuals was performed separately in each cohort under the additive model with SNPtest, v. 2.5.1. The analysis was additionally performed using untransformed apoA-I concentrations. Meta-analysis of the cohorts was performed with GWAMA software, v. 2.1, under the fixed-effect model. Genetic variants with $P<7x10^{-6}$ were considered as significant (Bonferroni-correction for 7,210 single nucleotide polymorphisms (SNPs) within 1-Mb region on either side flanking *APOA1*). Association of rs12225230 (with summary statistics available from a previous GWAS[10]) was replicated in our cohorts, and we therefore chose to use this SNP as an instrument in the Mendelian randomization analysis. Meta-analysis of the Finnish cohorts and the previous GWAS results[10] (total N=37,493) for rs12225230 was performed with GWAMA, v. 2.1. We estimated that the proportion of variance in apoA-I concentrations explained by rs12225230 was 0.0018.

We used the following steps to functionally annotate rs12225230: [1] To detect potential regulatory effects, we investigated colocalization with regulatory chromatin states and enhancer marks using the HaploReg database, v. 4.1[13]; and [2] To detect associations with DNA methylation in CpG sites, we investigated the mQTLdb[14] for methylation QTL (mQTL) effects. In addition, we investigated SNP-disease and SNP-trait associations from publicly available GWAS data using PhenoScanner v2[15], a curated database of human genotype-phenotype associations. Of note, when we conditioned on rs12225230, there were no significant association signals (Supplemental Figure 2), indicating that rs12225230 or SNPs in LD with this SNP are likely the only drivers of apoA-I association in this locus. Rs12225230 is a palindromic SNP (i.e., alleles on forward strand correspond to those on the reverse strand; G/C being C/G on the reverse strand); therefore, we additionally verified that the direction of effect used in the Mendelian randomization was correct by investigating the CAD associations of correlated ($r^2 \geq 0.85$), non-palindromic SNPs associated with apoA-I (e.g., rs11216162, rs78857232, rs76523703).

**Phenotype associations of rs12225230**

To investigate whether the association of rs12225230 with apoA-I level could be confounded by associations with other phenotypes, we screened publicly available genotype-phenotype associations using PhenoScanner.[15] The only significant associations using the traditional genome-wide significance threshold ($P<5x10^{-8}$) for rs12225230 were for apoA-I,[9, 10] HDL-C,[10, 16, 17] and total cholesterol levels.[16, 17] In addition, rs12225230 associated with height,[18] immature fraction of reticulocytes,[19] and triglyceride levels[20] ($P<1.7x10^{-5}$ using Bonferroni correction for 2,991 phenotypes analyzed). Given the potential association with triglycerides identified (P=3.2E-07 in a gene-centric study[21]), and the seemingly null association in our data (beta 0.001 for serum triglycerides, P=0.96), we performed a look-up of the variant in the largest GWAS to date of triglycerides (N up to 177,000)[17]: the association of rs12225230 with triglycerides was diluted close to null (beta 0.018, SE 0.0064, $p$ = 0.004). We generated a regional association plot (Supplemental Figure 4) of the triglyceride associations in this locus. We found that the triglyceride signal is separate from the apoA-I associations in this locus and happens to coincide in close proximity due to another apolipoprotein, possibly ApoC3 (the strongest signal for triglycerides being rs10790162, beta 0.231, SE 0.007, P=1.1E-249). Thus, rs12225230 was not associated with conventional vascular risk factors including type 2 diabetes, systolic blood pressure, BMI, smoking, or alcohol consumption (Supplemental Table 7).

**Additional data and data acknowledgements**

We used summary-level data from the following consortia/projects: meta-analysis of UK Biobank with CARDIoGRAMplusC4D,[22] meta-analysis of UK Biobank SOFT CAD GWAS (interim release) with CARDIoGRAMplusC4D 1000 Genomes-based GWAS and the Myocardial Infarction Genetics and CARDIoGRAM Exome,[23] Genotype-Tissue Expression Project (GTEx),[24] and MEGASTROKE.[25] Data on coronary artery disease and myocardial infarction have been contributed by the CARDIoGRAMplusC4D and UK Biobank CardioMetabolic Consortium CHD working group who used the UK Biobank Resource (application number 9922); data have been downloaded from www.CARDIOGRAMPLUSC4D.ORG. The Genotype-Tissue Expression (GTEx) Project was supported by the Common Fund of the Office of the Director of the National Institutes of Health, and by NCI, NHGRI, NHLBI, NIDA, NIMH, and NINDS; the data used for the analyses described in this manuscript were obtained from the GTEx Portal on 10/12/18. The MEGASTROKE project received funding from sources specified at http://www.megastroke.org/acknowledgments.html. The following authors contributed to the MEGASTROKE project: Rainer Malik, Ganesh Chauhan, Matthew Traylor, Muralidharan Sargurupremraj, Yukinori Okada, Aniket Mishra, Loes Rutten-Jacobs, Anne-Katrin Giese, Sander W van der Laan, Solveig Gretarsdottir, Christopher D Anderson,

Michael Chong, Hieab HH Adams, Tetsuro Ago, Peter Almgren, Philippe Amouyel, Hakan Ay, Traci M Bartz, Oscar R Benavente, Steve Bevan, Giorgio B Boncoraglio, Robert D Brown, Jr. , Adam S Butterworth, Caty Carrera, Cara L Carty, Daniel I Chasman, Wei-Min Chen, John W Cole, Adolfo Correa, Ioana Cotlarciuc, Carlos Cruchaga, John Danesh, Paul IW de Bakker, Anita L DeStefano, Marcel den Hoed, Qing Duan, Stefan T Engelter, Guido J Falcone, Rebecca F Gottesman, Raji P Grewal, Vilmundur Gudnason, Stefan Gustafsson, Jeffrey Haessler, Tamara B Harris, Ahamad Hassan, Aki S Havulinna, Susan R Heckbert, Elizabeth G Holliday, George Howard, Fang-Chi Hsu, Hyacinth I Hyacinth, M Arfan Ikram, Erik Ingelsson, Marguerite R Irvin, Xueqiu Jian, Jordi Jiménez-Conde, Julie A Johnson, J Wouter Jukema, Masahiro Kanai, Keith L Keene, Brett M Kissela, Dawn O Kleindorfer, Charles Kooperberg, Michiaki Kubo, Leslie A Lange, Carl D Langefeld, Claudia Langenberg, Lenore J Launer, Jin-Moo Lee, Robin Lemmens, Didier Leys, Cathryn M Lewis, Wei-Yu Lin, Arne G Lindgren, Erik Lorentzen, Patrik K Magnusson, Jane Maguire, Ani Manichaikul, Patrick F McArdle, James F Meschia, Braxton D Mitchell, Thomas H Mosley, Michael A Nalls, Toshiharu Ninomiya, Martin J O'Donnell, Bruce M Psaty, Sara L Pulit, Kristiina Rannikmäe, Alexander P Reiner, Kathryn M Rexrode, Kenneth Rice, Stephen S Rich, Paul M Ridker, Natalia S Rost, Peter M Rothwell, Jerome I Rotter, Tatjana Rundek, Ralph L Sacco, Saori Sakaue, Michele M Sale, Veikko Salomaa, Bishwa R Sapkota, Reinhold Schmidt, Carsten O Schmidt , Ulf Schminke, Pankaj Sharma, Agnieszka Slowik, Cathie LM Sudlow, Christian Tanislav, Turgut Tatlisumak, Kent D Taylor, Vincent NS Thijs, Gudmar Thorleifsson, Unnur Thorsteinsdottir, Steffen Tiedt, Stella Trompet, Christophe Tzourio, Cornelia M van Duijn, Matthew Walters, Nicholas J Wareham, Sylvia Wassertheil-Smoller, James G Wilson, Kerri L Wiggins, Qiong Yang, Salim Yusuf, Najaf Amin, Hugo S Aparicio, Donna K Arnett, John Attia, Alexa S Beiser, Claudine Berr, Julie E Buring, Mariana Bustamante, Valeria Caso, Yu-Ching Cheng, Seung Hoan Choi, Ayesha Chowhan, Natalia Cullell, Jean-François Dartigues, Hossein Delavaran, Pilar Delgado, Marcus Dörr, Gunnar Engström, Ian Ford, Wander S Gurpreet, Anders Hamsten, Laura Heitsch, Atsushi Hozawa, Laura Ibanez, Andreea Ilinca, Martin Ingelsson, Motoki Iwasaki, Rebecca D Jackson, Katarina Jood, Pekka Jousilahti, Sara Kaffashian, Lalit Kalra, Masahiro Kamouchi, Takanari Kitazono, Olafur Kjartansson, Manja Kloss, Peter J Koudstaal, Jerzy Krupinski, Daniel L Labovitz, Cathy C Laurie, Christopher R Levi, Linxin Li, Lars Lind, Cecilia M Lindgren, Vasileios Lioutas, Yong Mei Liu, Oscar L Lopez, Hirata Makoto, Nicolas Martinez-Majander, Koichi Matsuda, Naoko Minegishi, Joan Montaner , Andrew P Morris, Elena Muiño, Martina Müller-Nurasyid, Bo Norrving, Soichi Ogishima, Eugenio A Parati, Leema Reddy Peddareddygari, Nancy L Pedersen, Joanna Pera, Markus Perola, Alessandro Pezzini, Silvana

Pileggi, Raquel Rabionet, Iolanda Riba-Llena, Marta Ribasés, Jose R Romero, Jaume Roquer, Anthony G Rudd, Antti-Pekka Sarin, Ralhan Sarju, Chloe Sarnowski, Makoto Sasaki, Claudia L Satizabal, Mamoru Satoh, Naveed Sattar, Norie Sawada, Gerli Sibolt, Ásgeir Sigurdsson, Albert Smith, Kenji Sobue, Carolina Soriano-Tárraga, Tara Stanne, O Colin Stine, David J Stott, Konstantin Strauch, Takako Takai, Hideo Tanaka, Kozo Tanno, Alexander Teumer, Liisa Tomppo, Nuria P Torres-Aguila, Emmanuel Touze, Shoichiro Tsugane , Andre G Uitterlinden, Einar M Valdimarsson, Sven J van der Lee, Henry Völzke, Kenji Wakai , David Weir, Stephen R Williams, Charles DA Wolfe, Quenna Wong, Huichun Xu, Taiki Yamaji, Dharambir K Sanghera, Olle Melander, Christina Jern, Daniel Strbian, Israel Fernandez-Cadenas, W T Longstreth, Jr, Arndt Rolfs, Jun Hata, Daniel Woo, Jonathan Rosand, Guillaume Pare, Jemma C Hopewell, Danish Saleheen, Kari Stefansson, Bradford B Worrall, Steven J Kittner, Sudha Seshadri, Myriam Fornage, Hugh S Markus, Joanna MM Howson, Yoichiro Kamatani, Stephanie Debette, Martin Dichgans.

### Supplemental Figures

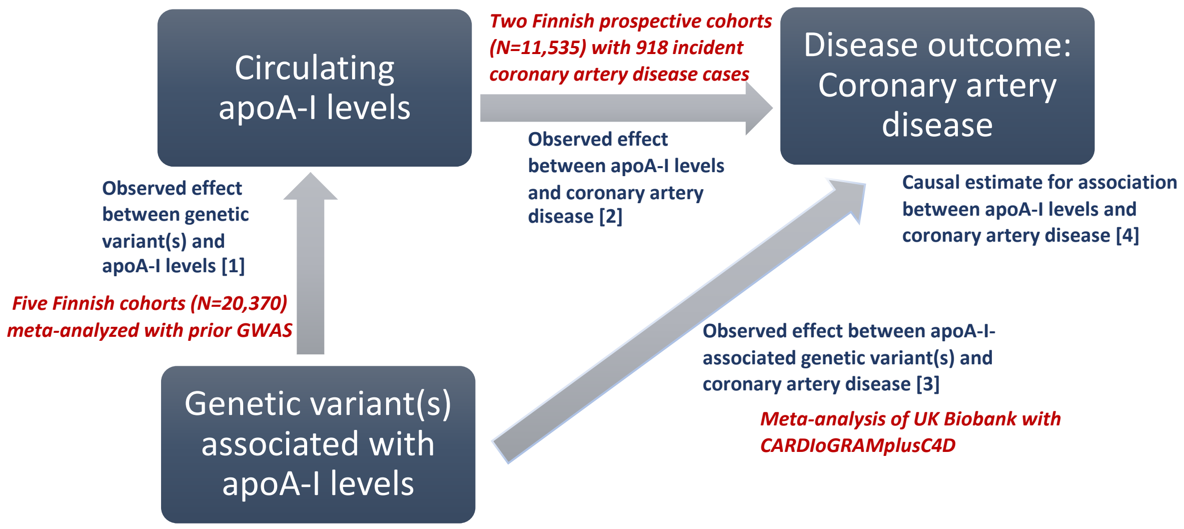

**Supplemental Figure 1. Flowchart showing the approach to estimate causality of circulating apoA-I levels in coronary artery disease.** Observed effects (between SNP and apoA-I [1], and between SNP and CAD [3]) can be used to calculate a causal estimate for apoA-I-CAD association [4]. This estimate is compared to the observed apoA-I-CAD [2] association. If the estimates are of a similar magnitude, apoA-I levels would be causal (protective) for CAD. The data sources used in the present study are shown. Data from five Finnish cohorts were used. To obtain an estimate for SNP-apoA-I association [1], these data were meta-analyzed with previous GWAS of apoA-I[10]. SNP-CAD association [3] was obtained from a large meta-analysis of CAD[22] as indicated.

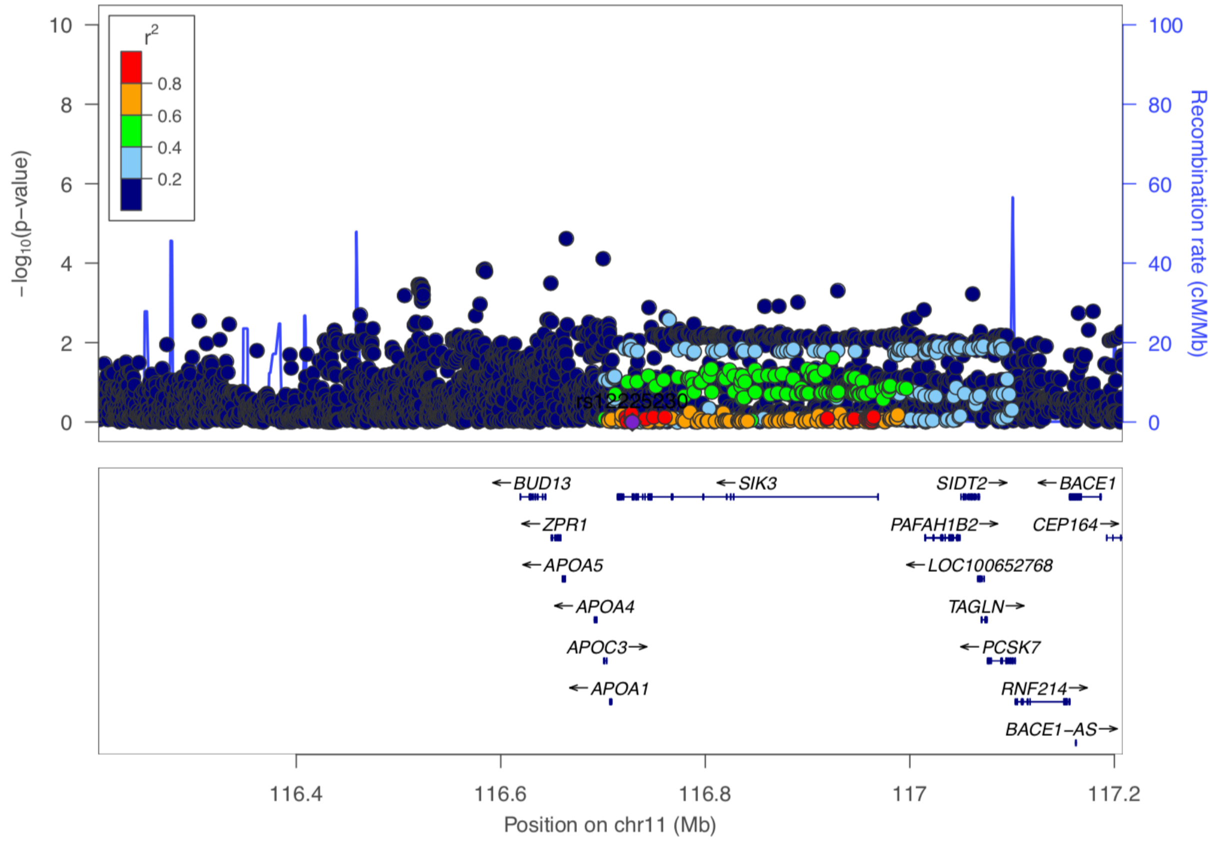

**Supplemental Figure 2. Regional association plot showing the association of *APOA1* locus SNPs with circulating apoA-I levels when conditioning on rs12225230.** There were no association signals in the *APOA1* locus when rs12225230 (SNP highlighted in violet) was conditioned on in the association analysis. This indicates that rs12225230 and SNPs in LD with this SNP are the only drivers of association in this locus.

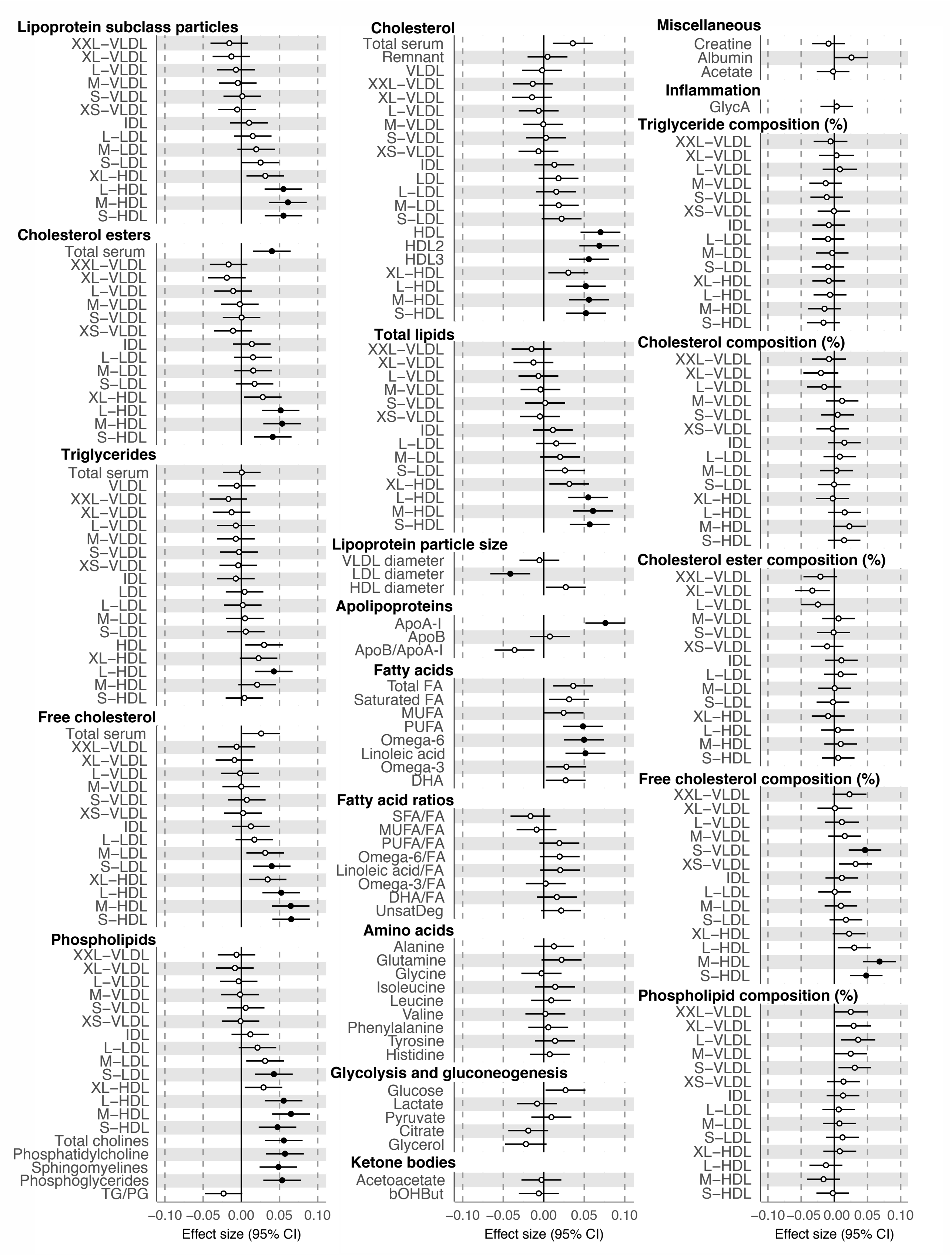

**Supplemental Figure 3. Forest plot showing association of rs12225230 with all the metabolic measures from the serum metabolomics experiments.** Effect sizes (betas) are given as standard deviation (SD) units in inverse-rank transformed covariate-adjusted metabolite residuals. Closed symbols, P<0.002 (significant association after Bonferroni correction with 22 principal components that explained >95% of total variation in the levels of the metabolic measures); open symbols P>0.002. See Supplemental Table 4 for full description of the metabolic measures.

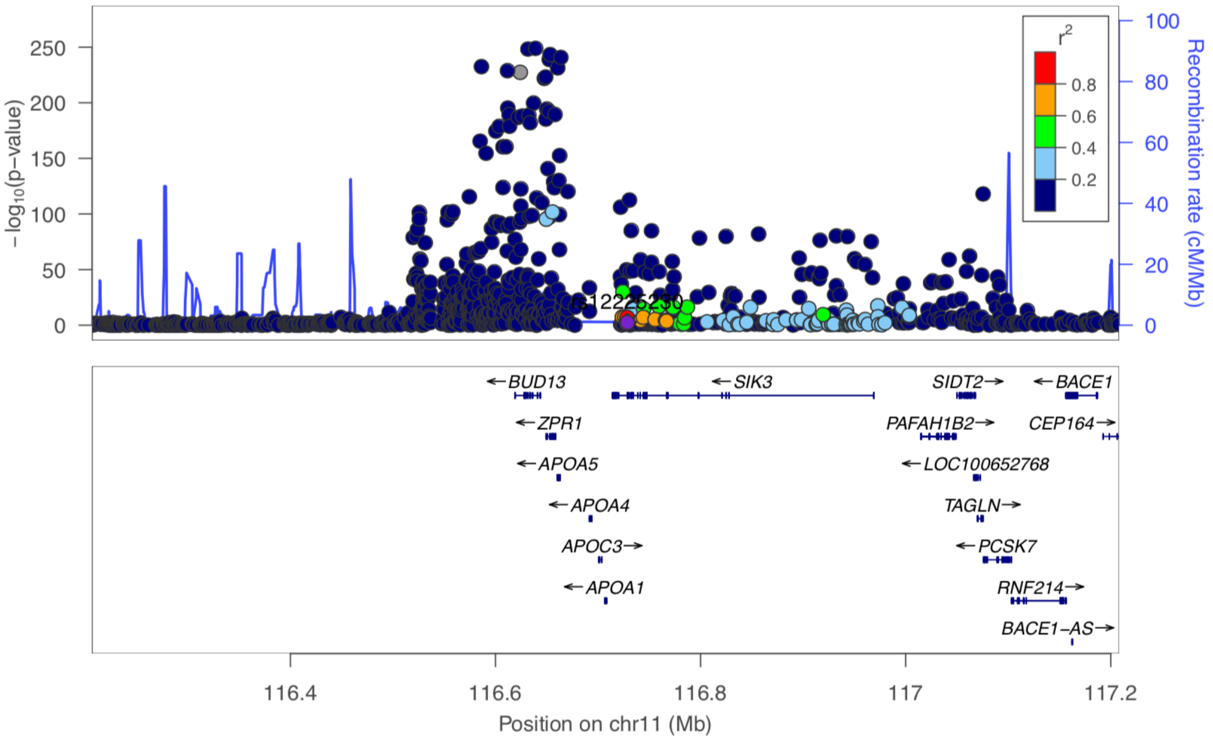

**Supplemental Figure 4. Regional association plot showing the association of *APOA1* locus SNPs with circulating total triglycerides.** The triglyceride associations are from a large GWAS of circulating lipid levels.[17] The strongest signal for triglycerides was rs10790162 (beta 0.231, SE 0.007, P=1.1E-249). Rs12225230 that was used as an instrument here (SNP highlighted in violet) was not associated with triglyceride levels (and was not in LD with rs10790162; $r^2$<0.01).

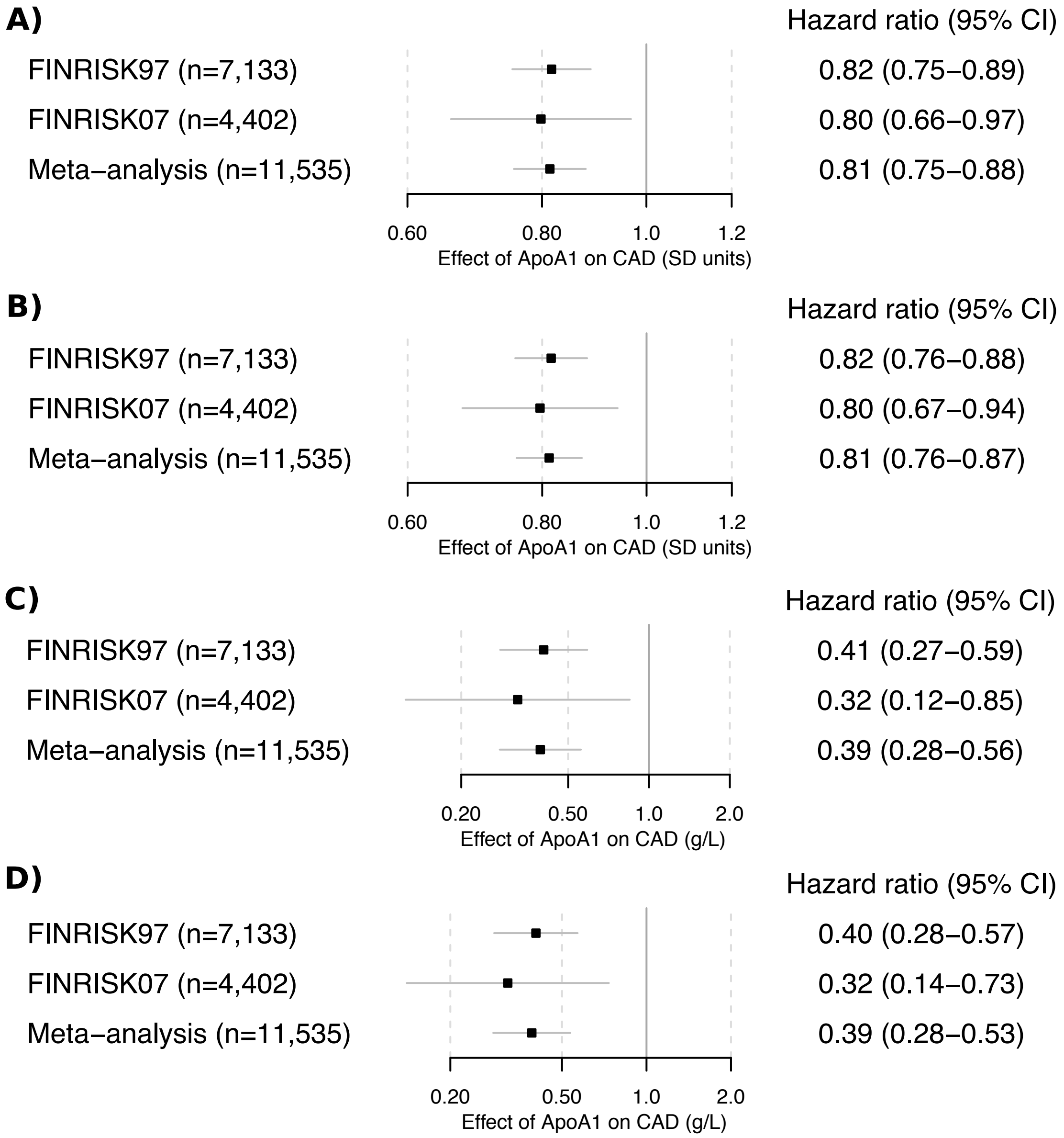

**Supplemental Figure 5. Forest plots showing observational associations of circulating apoA-I levels with coronary artery disease.** Observational associations were estimated in the FINRISK97 and FINRISK07 cohorts under two models and are presented *per* SD unit (A, B) or *per* 1 g/L increase in apoA-I (C, D)**:** (A, C) Fully-adjusted model (age, sex, LDL cholesterol, body-mass index, systolic blood pressure, type 2 diabetes, smoking and alcohol consumption as covariates); (B, D) Minimaly-adjusted model (age and sex as covariates). The effect estimates were consistent in both cohorts and under full and minimal models.

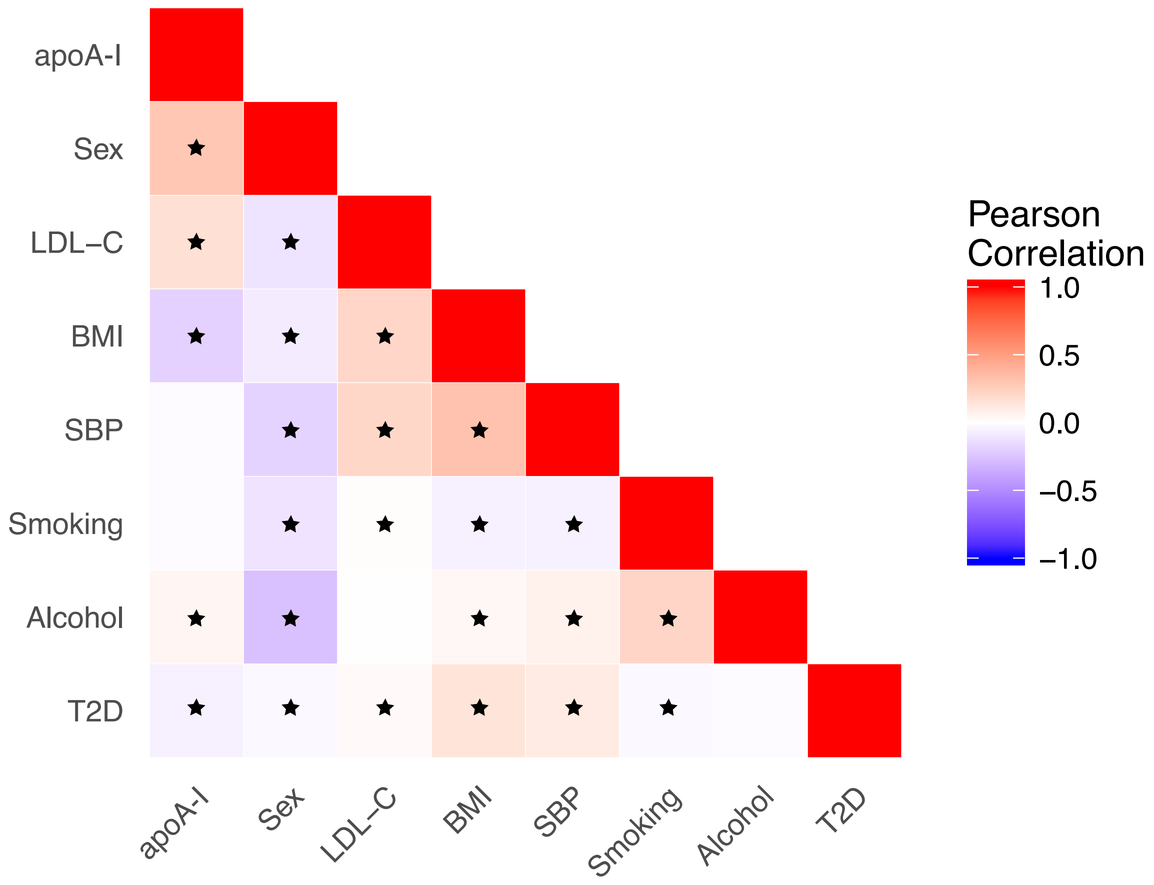

**Supplemental Figure 6. Heatmap showing correlations between circulating apoA-I levels and covariates included in the regression models.** Correlation of circulating apoA-I levels with potential confounders (covariates included in the model when calculating the observational estimate for apoA-I-CAD association) was estimated in the FINRISK97 cohort. Significant (P<0.008 using Bonferroni correction for 28 pairwise comparisons) correlations are indicated by an asterisk. Correlations of apoA-I with the covariates were low (*r* between -0.3 and 0.3); highest positive and negative correlations were with sex (*r*=0.29) and BMI (*r*=-0.19), respectively. Abbreviations: LDL-C, LDL-cholesterol; BMI, body mass index; SBP, systolic blood pressure; smoking, current smoking; alcohol, alcohol consumption; T2D, type 2 diabetes.

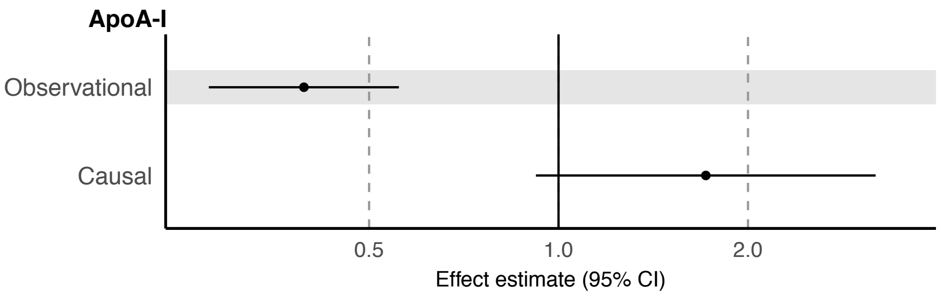

**Supplemental Figure 7. Observational and causal effects of circulating apoA-I levels and risk of coronary artery disease.** Observational estimate was assessed in the FINRISK97 and FINRISK07 cohorts with age, sex, LDL cholesterol, body-mass index, systolic blood pressure, type 2 diabetes, smoking and alcohol consumption as covariates. Causal estimate was calculated based on association of rs12225230 (associated with increased serum apoA-I) with coronary artery disease (CAD) in meta-analysis of the UK Biobank and CARDIoGRAMplusC4D data[23]. The estimates were calculated for 1 unit (g/L) increase in apoA-I concentration (observational estimate, HR 0.39, 95%CI 0.28-0.56; causal estimate, OR 1.71, 95%CI 0.92-3.19). The MR estimate was scaled to SD units (shown in Figure 4).

### Supplemental Tables

#### Supplemental Table 1. Characteristics of the study populations used in the genetic analyses.

| Characteristic | FINRISK97 | FINRISK07 | NFBC66 | NFBC86 | YFS |
|---|---|---|---|---|---|
| Number of participants | 6,643 | 3,896 | 4,668 | 3,215 | 1,948 |
| Age (year ± SD) | 47.7 ± 13.1 | 50.3 ± 13.4 | 31.0 ± 0.3 | 16.1 ± 0.4 | 37.7 ± 5.0 |
| Female (%) | 52 | 55 | 52 | 51 | 54 |

#### Supplemental Table 2. Characteristics of the study populations used in the observational analyses.

| Characteristic | FINRISK97 | FINRISK07 |
|---|---|---|
| Number of participants | 7,133 | 4,402 |
| Age (year ± SD) | 47.8. ± 13.0 | 51.8 ± 13.4 |
| Female (%) | 50 | 53 |
| Incident coronary artery disease cases, *n* (%) | 743 (10) | 175 (4) |
| apoA-I, g/L (mean ± SD) | 1.579 ± 0.225 | 1.533 ± 0.200 |
| LDL cholesterol, mmol/L (mean ± SD) | 1.9 ± 0.6 | 1.6 ± 0.5 |
| Body mass index (mean ± SD) | 26.6 ± 4.5 | 27.1 ± 4.7 |
| Systolic blood pressure (mean ± SD) | 136 ± 20 | 137 ± 20 |
| Type 2 diabetes cases, *n* (%) | 384 (5) | 360 (8) |
| Smoking (%) | 24 | 18 |
| Alcohol use, g / week [median (range)] | 26 (0-2,439) | 25 (0-1,590) |

#### Supplemental Table 3. Power considerations for the Mendelian randomization study.

| Effect estimate (odds ratio)[a] | Power[b] |
|---|---|
| 0.70 | 0.99 |
| 0.80 | 0.78 |
| 0.90 | 0.27 |

[a] Odds ratio for the effect of circulating apoA-I levels with risk of coronary artery disease (*per* SD unit higher apoA-I) .
[b]Power was assessed with mRnd (http://cnsgenomics.com/shiny/mRnd/) using rs12225230 as the genetic instrument (at a 2-sided alpha of 0.05 and sample size of 547,261 totaling 122,733 CAD cases[22]).

**Supplemental Table 4. List of metabolic measures.**

| Metabolic measure | Description |
|---|---|
| **Lipoprotein subclass particle concentrations** | |
| Extremely Large VLDL (XXL-VLDL) | Concentration of chylomicrons and extremely large VLDL particles |
| Very Large VLDL (XL-VLDL) | Concentration of very large VLDL particles |
| Large VLDL (L-VLDL) | Concentration of large VLDL particles |
| Medium VLDL (M-VLDL) | Concentration of medium VLDL particles |
| Small VLDL (S-VLDL) | Concentration of small VLDL particles |
| Very Small VLDL (XS-VLDL) | Concentration of very small VLDL particles |
| IDL | Concentration of IDL particles |
| Large LDL (L-LDL) | Concentration of large LDL particles |
| Medium LDL (M-LDL) | Concentration of medium LDL particles |
| Small LDL (S-LDL) | Concentration of small LDL particles |
| Very Large HDL (XL-HDL) | Concentration of very large HDL particles |
| Large HDL (L-HDL) | Concentration of large HDL particles |
| Medium HDL (M-HDL) | Concentration of medium HDL particles |
| Small HDL (S-HDL) | Concentration of small HDL particles |
| **Cholesterol** | |
| Total serum cholesterol | Serum total cholesterol |
| Remnant | Remnant cholesterol (non-HDL, non-LDL -cholesterol) |
| VLDL | Total cholesterol in VLDL |
| Extremely Large VLDL (XXL-VLDL) | Total cholesterol in chylomicrons and extremely large VLDL |
| Very Large VLDL (XL-VLDL) | Total cholesterol in very large VLDL |
| Large VLDL (L-VLDL) | Total cholesterol in large VLDL |
| Medium VLDL (M-VLDL) | Total cholesterol in medium VLDL |
| Small VLDL (S-VLDL) | Total cholesterol in small VLDL |
| Very small VLDL (XS-VLDL) | Total cholesterol in very small VLDL |
| IDL | Total cholesterol in IDL |
| LDL | Total cholesterol in LDL |
| Large LDL (L-LDL) | Total cholesterol in large LDL |
| Medium LDL (M-LDL) | Total cholesterol in medium LDL |
| Small LDL (S-LDL) | Total cholesterol in small LDL |
| HDL | Total cholesterol in HDL |
| HDL2 | Total cholesterol in HDL2 |
| HDL3 | Total cholesterol in HDL3 |
| Very Large HDL (XL-HDL) | Total cholesterol in very large HDL |
| Large HDL (L-HDL) | Total cholesterol in large HDL |
| Medium HDL (M-HDL) | Total cholesterol in medium HDL |
| Small HDL (S-HDL) | Total cholesterol in small HDL |
| **Triglycerides** | |
| Total serum triglycerides (TG) | Serum total triglycerides |
| VLDL | Triglycerides in VLDL |
| Extremely Large VLDL (XXL-VLDL) | Triglycerides in chylomicrons and extremely large VLDL |
| Very Large VLDL (XL-VLDL) | Triglycerides in very large VLDL |
| Large VLDL (L-VLDL) | Triglycerides in large VLDL |
| Medium VLDL (M-VLDL) | Triglycerides in medium VLDL |
| Small VLDL (S-VLDL) | Triglycerides in small VLDL |
| Very small VLDL (XS-VLDL) | Triglycerides in very small VLDL |
| IDL | Triglycerides in IDL |
| LDL | Triglycerides in LDL |
| Large LDL (L-LDL) | Triglycerides in large LDL |
| Medium LDL (M-LDL) | Triglycerides in medium LDL |
| Small LDL (S-LDL) | Triglycerides in small LDL |
| HDL | Triglycerides in HDL |
| Very Large HDL (XL-HDL) | Triglycerides in very large HDL |

| Metabolic measure | Description |
|---|---|
| Large HDL (L-HDL) | Triglycerides in large HDL |
| Medium HDL (M-HDL) | Triglycerides in medium HDL |
| Small HDL (S-HDL) | Triglycerides in small HDL |
| **Phospholipids** | |
| Extremely Large VLDL (XXL-VLDL) | Phospholipids in chylomicrons and extremely large VLDL |
| Very Large VLDL (XL-VLDL) | Phospholipids in very large VLDL |
| Large VLDL (L-VLDL) | Phospholipids in large VLDL |
| Medium VLDL (M-VLDL) | Phospholipids in medium VLDL |
| Small VLDL (S-VLDL) | Phospholipids in small VLDL |
| Very small VLDL (XS-VLDL) | Phospholipids in very small VLDL |
| IDL | Phospholipids in IDL |
| Large LDL (L-LDL) | Phospholipids in large LDL |
| Medium LDL (M-LDL) | Phospholipids in medium LDL |
| Small LDL (S-LDL) | Phospholipids in small LDL |
| Very Large HDL (XL-HDL) | Phospholipids in very large HDL |
| Large HDL (L-HDL) | Phospholipids in large HDL |
| Medium HDL (M-HDL) | Phospholipids in medium HDL |
| Small HDL (S-HDL) | Phospholipids in small HDL |
| Cholines | Total cholines |
| Phosphatidylcholine | Phosphatidylcholine and other cholines |
| Sphingomyelins | Sphingomyelins |
| Phosphoglycerides | Total phosphoglycerides |
| TG to PG ratio (%) (TG/PG) | Ratio of triglycerides to phosphoglycerides |
| **Total lipids** | |
| Extremely Large VLDL (XXL-VLDL) | Total lipids in chylomicrons and extremely large VLDL |
| Very Large VLDL (XL-VLDL) | Total lipids in very large VLDL |
| Large VLDL (L-VLDL) | Total lipids in large VLDL |
| Medium VLDL (M-VLDL) | Total lipids in medium VLDL |
| Small VLDL (S-VLDL) | Total lipids in small VLDL |
| Very small VLDL (XS-VLDL) | Total lipids in very small VLDL |
| IDL | Total lipids in IDL |
| Large LDL (L-LDL) | Total lipids in large LDL |
| Medium LDL (M-LDL) | Total lipids in medium LDL |
| Small LDL (S-LDL) | Total lipids in small LDL |
| Very Large HDL (XL-HDL) | Total lipids in very large HDL |
| Large HDL (L-HDL) | Total lipids in large HDL |
| Medium HDL (M-HDL) | Total lipids in medium HDL |
| Small HDL (S-HDL) | Total lipids in small HDL |
| **Cholesterol esters** | |
| Esterified cholesterol, total serum | Esterified cholesterol |
| Extremely Large VLDL (XXL-VLDL) | Cholesterol esters in chylomicrons and extremely large VLDL |
| Very Large VLDL (XL-VLDL) | Cholesterol esters in very large VLDL |
| Large VLDL (L-VLDL) | Cholesterol esters in large VLDL |
| Medium VLDL (M-VLDL) | Cholesterol esters in medium VLDL |
| Small VLDL (S-VLDL) | Cholesterol esters in small VLDL |
| Very small VLDL (XS-VLDL) | Cholesterol esters in very small VLDL |
| IDL | Cholesterol esters in IDL |
| Large LDL (L-LDL) | Cholesterol esters in large LDL |
| Medium LDL (M-LDL) | Cholesterol esters in medium LDL |
| Small LDL (S-LDL) | Cholesterol esters in small LDL |
| Very Large HDL (XL-HDL) | Cholesterol esters in very large HDL |
| Large HDL (L-HDL) | Cholesterol esters in large HDL |
| Medium HDL (M-HDL) | Cholesterol esters in medium HDL |
| Small HDL (S-HDL) | Cholesterol esters in small HDL |
| **Free cholesterol** | |

| Metabolic measure | Description |
|---|---|
| Free cholesterol, total serum | Free cholesterol |
| Extremely Large VLDL (XXL-VLDL) | Free cholesterol in chylomicrons and extremely large VLDL |
| Very Large VLDL (XL-VLDL) | Free cholesterol in very large VLDL |
| Large VLDL (L-VLDL) | Free cholesterol in large VLDL |
| Medium VLDL (M-VLDL) | Free cholesterol in medium VLDL |
| Small VLDL (S-VLDL) | Free cholesterol in small VLDL |
| Very small VLDL (XS-VLDL) | Free cholesterol in very small VLDL |
| IDL | Free cholesterol in IDL |
| Large LDL (L-LDL) | Free cholesterol in large LDL |
| Medium LDL (M-LDL) | Free cholesterol in medium LDL |
| Small LDL (S-LDL) | Free cholesterol in small LDL |
| Very Large HDL (XL-HDL) | Free cholesterol in very large HDL |
| Large HDL (L-HDL) | Free cholesterol in large HDL |
| Medium HDL (M-HDL) | Free cholesterol in medium HDL |
| Small HDL (S-HDL) | Free cholesterol in small HDL |
| **Lipoprotein particle size** | |
| VLDL particle size (VLDL diameter) | Mean diameter for VLDL particles |
| LDL particle size (LDL diameter) | Mean diameter for LDL particles |
| HDL particle size (HDL diameter) | Mean diameter for HDL particles |
| **Apolipoproteins** | |
| Apolipoprotein A-I (ApoA-I) | Apolipoprotein A-I |
| Apolipoprotein B (ApoB) | Apolipoprotein B |
| ApoB to ApoAI ratio (ApoB/ApoA-I) | Ratio of apolipoprotein B to apolipoprotein A-I |
| **Fatty acids** | |
| Total fatty acids (FA) | Total fatty acids |
| Saturated fatty acids (FA) | Saturated fatty acids |
| MUFA | Monounsaturated fatty acids; 16:1, 18:1 |
| PUFA | Polyunsaturated fatty acids |
| Omega-6 | Omega-6 fatty acids |
| Linoleic acid | 18:2, linoleic acid |
| Omega-3 | Omega-3 fatty acids |
| Docosahexaenoic acid (DHA) | 22:6, docosahexaenoic acid |
| **Fatty acid ratios** | |
| Saturated fatty acids (%) (SFA/FA) | Ratio of saturated fatty acids to total fatty acids |
| MUFA (%) (MUFA/FA) | Ratio of monounsaturated fatty acids to total fatty acids |
| PUFA (%) (PUFA/FA) | Ratio of polyunsaturated fatty acids to total fatty acids |
| Omega-6 (%) (Omega-6/FA) | Ratio of omega-6 fatty acids to total fatty acids |
| Linoleic acid (%) (Linoleic acid/FA) | Ratio of 18:2 linoleic acid to total fatty acids |
| Omega-3 (%) (Omega-3/FA) | Ratio of omega-3 fatty acids to total fatty acids |
| Docosahexaenoic acid (%) (DHA/FA) | Ratio of 22:6 docosahexaenoic acid to total fatty acids |
| Degree of unsaturation (UnSatDeg) | Estimated degree of unsaturation |
| **Amino acids** | |
| Alanine | Alanine |
| Glutamine | Glutamine |
| Glycine | Glycine |
| Isoleucine | Isoleucine |
| Leucine | Leucine |
| Valine | Valine |
| Phenylalanine | Phenylalanine |
| Tyrosine | Tyrosine |
| Histidine | Histidine |
| **Glycolysis and gluconeogenesis** | |
| Glucose | Glucose |
| Lactate | Lactate |
| Pyruvate | Pyruvate |

| Metabolic measure | Description |
|---|---|
| Citrate | Citrate |
| Glycerol | Glycerol |
| **Ketone bodies** | |
| Acetoacetate | Acetoacetate |
| Beta-hydroxybutyrate (bOHBut) | 3-hydroxybutyrate |
| **Miscellaneous** | |
| Creatinine | Creatinine |
| Albumin | Albumin |
| Acetate | Acetate |
| **Inflammation** | |
| Glycoprotein acetyls (GlycA) | Glycoprotein acetyls, mainly a1-acid glycoprotein |
| **Cholesterol composition (%)** | |
| Extremely Large VLDL (XXL-VLDL) | Total cholesterol to total lipids ratio in chylomicrons and extremely large VLDL |
| Very Large VLDL (XL-VLDL) | Total cholesterol to total lipids ratio in very large VLDL |
| Large VLDL (L-VLDL) | Total cholesterol to total lipids ratio in large VLDL |
| Medium VLDL (M-VLDL) | Total cholesterol to total lipids ratio in medium VLDL |
| Small VLDL (S-VLDL) | Total cholesterol to total lipids ratio in small VLDL |
| Very small VLDL (XS-VLDL) | Total cholesterol to total lipids ratio in very small VLDL |
| IDL | Total cholesterol to total lipids ratio in IDL |
| Large LDL (L-LDL) | Total cholesterol to total lipids ratio in large LDL |
| Medium LDL (M-LDL) | Total cholesterol to total lipids ratio in medium LDL |
| Small LDL (S-LDL) | Total cholesterol to total lipids ratio in small LDL |
| Very Large HDL (XL-HDL) | Total cholesterol to total lipids ratio in very large HDL |
| Large HDL (L-HDL) | Total cholesterol to total lipids ratio in large HDL |
| Medium HDL (M-HDL) | Total cholesterol to total lipids ratio in medium HDL |
| Small HDL (S-HDL) | Total cholesterol to total lipids ratio in small HDL |
| **Triglyceride composition (%)** | |
| Extremely Large VLDL (XXL-VLDL) | Triglycerides to total lipids ratio in chylomicrons and extremely large VLDL |
| Very Large VLDL (XL-VLDL) | Triglycerides to total lipids ratio in very large VLDL |
| Large VLDL (L-VLDL) | Triglycerides to total lipids ratio in large VLDL |
| Medium VLDL (M-VLDL) | Triglycerides to total lipids ratio in medium VLDL |
| Small VLDL (S-VLDL) | Triglycerides to total lipids ratio in small VLDL |
| Very small VLDL (XS-VLDL) | Triglycerides to total lipids ratio in very small VLDL |
| IDL | Triglycerides to total lipids ratio in IDL |
| Large LDL (L-LDL) | Triglycerides to total lipids ratio in large LDL |
| Medium LDL (M-LDL) | Triglycerides to total lipids ratio in medium LDL |
| Small LDL (S-LDL) | Triglycerides to total lipids ratio in small LDL |
| Very Large HDL (XL-HDL) | Triglycerides to total lipids ratio in very large HDL |
| Large HDL (L-HDL) | Triglycerides to total lipids ratio in large HDL |
| Medium HDL (M-HDL) | Triglycerides to total lipids ratio in medium HDL |
| Small HDL (S-HDL) | Triglycerides to total lipids ratio in small HDL |
| **Cholesterol ester composition (%)** | |
| Extremely Large VLDL (XXL-VLDL) | Cholesterol esters to total lipids ratio in chylomicrons and extremely large VLDL |
| Very Large VLDL (XL-VLDL) | Cholesterol esters to total lipids ratio in very large VLDL |
| Large VLDL (L-VLDL) | Cholesterol esters to total lipids ratio in large VLDL |
| Medium VLDL (M-VLDL) | Cholesterol esters to total lipids ratio in medium VLDL |
| Small VLDL (S-VLDL) | Cholesterol esters to total lipids ratio in small VLDL |
| Very small VLDL (XS-VLDL) | Cholesterol esters to total lipids ratio in very small VLDL |
| IDL | Cholesterol esters to total lipids ratio in IDL |
| Large LDL (L-LDL) | Cholesterol esters to total lipids ratio in large LDL |
| Medium LDL (M-LDL) | Cholesterol esters to total lipids ratio in medium LDL |
| Small LDL (S-LDL) | Cholesterol esters to total lipids ratio in small LDL |

| Metabolic measure | Description |
|---|---|
| Very Large HDL (XL-HDL) | Cholesterol esters to total lipids ratio in very large HDL |
| Large HDL (L-HDL) | Cholesterol esters to total lipids ratio in large HDL |
| Medium HDL (M-HDL) | Cholesterol esters to total lipids ratio in medium HDL |
| Small HDL (S-HDL) | Cholesterol esters to total lipids ratio in small HDL |
| **Free cholesterol composition (%)** | |
| Extremely Large VLDL (XXL-VLDL) | Free cholesterol to total lipids ratio in chylomicrons and extremely large VLDL |
| Very Large VLDL (XL-VLDL) | Free cholesterol to total lipids ratio in very large VLDL |
| Large VLDL (L-VLDL) | Free cholesterol to total lipids ratio in large VLDL |
| Medium VLDL (M-VLDL) | Free cholesterol to total lipids ratio in medium VLDL |
| Small VLDL (S-VLDL) | Free cholesterol to total lipids ratio in small VLDL |
| Very small VLDL (XS-VLDL) | Free cholesterol to total lipids ratio in very small VLDL |
| IDL | Free cholesterol to total lipids ratio in IDL |
| Large LDL (L-LDL) | Free cholesterol to total lipids ratio in large LDL |
| Medium LDL (M-LDL) | Free cholesterol to total lipids ratio in medium LDL |
| Small LDL (S-LDL) | Free cholesterol to total lipids ratio in small LDL |
| Very Large HDL (XL-HDL) | Free cholesterol to total lipids ratio in very large HDL |
| Large HDL (L-HDL) | Free cholesterol to total lipids ratio in large HDL |
| Medium HDL (M-HDL) | Free cholesterol to total lipids ratio in medium HDL |
| Small HDL (S-HDL) | Free cholesterol to total lipids ratio in small HDL |
| **Phospholipid composition (%)** | |
| Extremely Large VLDL (XXL-VLDL) | Phospholipids to total lipids ratio in chylomicrons and extremely large VLDL |
| Very Large VLDL (XL-VLDL) | Phospholipids to total lipids ratio in very large VLDL |
| Large VLDL (L-VLDL) | Phospholipids to total lipids ratio in large VLDL |
| Medium VLDL (M-VLDL) | Phospholipids to total lipids ratio in medium VLDL |
| Small VLDL (S-VLDL) | Phospholipids to total lipids ratio in small VLDL |
| Very small VLDL (XS-VLDL) | Phospholipids to total lipids ratio in very small VLDL |
| IDL | Phospholipids to total lipids ratio in IDL |
| Large LDL (L-LDL) | Phospholipids to total lipids ratio in large LDL |
| Medium LDL (M-LDL) | Phospholipids to total lipids ratio in medium LDL |
| Small LDL (S-LDL) | Phospholipids to total lipids ratio in small LDL |
| Very Large HDL (XL-HDL) | Phospholipids to total lipids ratio in very large HDL |
| Large HDL (L-HDL) | Phospholipids to total lipids ratio in large HDL |
| Medium HDL (M-HDL) | Phospholipids to total lipids ratio in medium HDL |
| Small HDL (S-HDL) | Phospholipids to total lipids ratio in small HDL |

**Supplemental Table 5. Randomized clinical trials of apoA-I infusion products reporting cardiovascular end points.**

| Drug / apoA-I source[a] | Trial | Dose | Population recruitment and follow-up | End points | Reference |
|---|---|---|---|---|---|
| CSL111 / native plasma apoA-I | ERASE (NCT00225719) | 40 or 80 mg/kg weekly for 4 weeks | *Patients with clinical need for coronary angiography (n=183)<br>*July 2005-October 2006 | *Intravascular ultrasonography (IVUS) measurements | Tardif *et al.* 2007[26] |
| CER-001 / recombinant wild-type apoA-I | CHI-SQUARE, phase 2 (NCT01201837) | 3, 6 or 12 mg/kg weekly for 6 weeks | *Patients with acute coronary syndromes (n=507)<br>*March 2011-August 2012<br>*Follow-up 6 months after last infusion | *IVUS and quantitative coronary angiography measurements<br>***MACE end points (cardiac arrest, non-fatal MI, non-fatal stroke, coronary revascularization, unstable angina, heart failure)** | Tardif *et al.* 2014[27] |
| | CARAT, phase 2 (NCT02484378) | 3 mg/kg (~0.225 g) weekly for 10 weeks | *Patients with acute coronary syndromes (n=171)<br>*August 2015-November 2016<br>*Follow-up from baseline to day 78 (7-21 days after last infusion) | *IVUS measurements | Nicholls *et al.* 2018[28] |
| CSL112 / native plasma apoA-I | AEGIS-I, phase 2 (NCT02108262) | 2 or 6 g weekly for 4 weeks (6 g corresponds to ~80 mg/kg) | *Patients with acute myocardial infarction (n= 1,258)<br>*January-November 2015<br>*Follow-up: Up to 1 year after randomization (89 patients) or until the last randomized subject completed the study day 112 visit | ***MACE end points (cardiovascular death, nonfatal MI, ischemic stroke, unstable angina, all-cause mortality, non-cardiovascular death, hemorrhagic stroke, any stroke, heart failure, coronary revascularization)** | Gibson *et al.* 2016[29] |
| MDCO-216 / recombinant apoA-I-Milano | MILANO PILOT, phase 2 (NCT02678923) | 20 mg/kg (~1.5 g) weekly for 5 weeks | *Patients with acute coronary syndromes (n=122)<br>*December 2015-August 2016<br>*Follow-up at day 36 | *IVUS measurements | Nicholls *et al.* 2018[30] |

[a]Placebo-controlled phase II RCTs using apoA-I infusion products are shown. Two of the trials reported major adverse cardiovascular event (MACE) end points (in bold typeface above).

### Supplemental Table 6. Association of *APOA1* locus SNPs with circulating apoA-I levels.

| SNP[a] | Position on chromosome 11 | Reference allele | Other allele | Effect allele frequency | Beta[b] | Standar d error | p-value |
|---|---|---|---|---|---|---|---|
| rs186911592 | 116629483 | G | T | 0.053 | 0.111 | 0.022 | 5.03E-07 |
| rs140575350 | 116646820 | G | C | 0.070 | 0.098 | 0.019 | 5.34E-07 |
| rs182732109 | 116687695 | G | T | 0.051 | 0.117 | 0.023 | 2.34E-07 |
| rs5092 | 116693464 | T | C | 0.735 | -0.052 | 0.011 | 3.81E-06 |
| rs10892035 | 116694938 | G | A | 0.735 | -0.052 | 0.011 | 4.62E-06 |
| rs12721078 | 116699395 | A | C | 0.065 | 0.116 | 0.020 | 9.70E-09 |
| rs2854117 | 116700142 | C | T | 0.664 | -0.048 | 0.011 | 4.91E-06 |
| rs4520 | 116701535 | C | T | 0.634 | -0.051 | 0.010 | 9.33E-07 |
| rs2070666 | 116701674 | A | T | 0.202 | 0.057 | 0.012 | 4.41E-06 |
| rs11216153 | 116705100 | T | G | 0.219 | 0.060 | 0.012 | 7.30E-07 |
| rs12721030 | 116705278 | T | C | 0.238 | 0.059 | 0.012 | 4.73E-07 |
| rs525028 | 116705516 | A | G | 0.623 | -0.054 | 0.010 | 1.83E-07 |
| rs12721028 | 116705590 | G | A | 0.215 | 0.058 | 0.012 | 2.06E-06 |
| rs670 | 116708413 | T | C | 0.188 | 0.070 | 0.013 | 3.76E-08 |
| rs613808 | 116710968 | G | A | 0.621 | -0.048 | 0.010 | 2.44E-06 |
| rs61905145 | 116718415 | A | G | 0.178 | 0.070 | 0.013 | 8.42E-08 |
| rs1145206 | 116718521 | C | G | 0.188 | 0.074 | 0.013 | 6.23E-09 |
| rs111894427 | 116721405 | T | C | 0.065 | 0.116 | 0.020 | 9.91E-09 |
| rs688456 | 116722551 | T | G | 0.212 | 0.071 | 0.012 | 4.93E-09 |
| rs888245 | 116723737 | G | C | 0.178 | 0.071 | 0.013 | 3.56E-08 |
| rs12099358 | 116726048 | A | C | 0.182 | 0.070 | 0.013 | 4.04E-08 |
| rs61907560 | 116726087 | C | T | 0.065 | 0.115 | 0.020 | 1.16E-08 |
| rs61907561 | 116726796 | G | C | 0.063 | 0.118 | 0.020 | 7.94E-09 |
| rs78857232 | 116727465 | T | C | 0.170 | 0.078 | 0.013 | 2.70E-09 |
| rs61907562 | 116727592 | A | G | 0.065 | 0.115 | 0.020 | 1.19E-08 |
| rs625145 | 116727936 | T | A | 0.204 | 0.076 | 0.012 | 6.18E-10 |
| rs11216162 | 116728277 | A | G | 0.192 | 0.076 | 0.013 | 1.55E-09 |
| rs12225230 | 116728630 | C | G | 0.192 | 0.076 | 0.013 | 1.47E-09 |
| rs61907563 | 116733565 | A | G | 0.064 | 0.115 | 0.020 | 1.60E-08 |
| rs11216168 | 116741553 | A | G | 0.171 | 0.077 | 0.013 | 4.24E-09 |
| rs17120111 | 116744018 | T | C | 0.183 | 0.068 | 0.013 | 9.51E-08 |
| rs76523703 | 116749509 | G | T | 0.169 | 0.078 | 0.013 | 3.02E-09 |
| rs11827828 | 116755584 | A | G | 0.177 | 0.072 | 0.013 | 2.41E-08 |
| rs10892042 | 116760545 | A | G | 0.168 | 0.078 | 0.013 | 3.64E-09 |
| rs11820814 | 116765700 | C | T | 0.169 | 0.077 | 0.013 | 5.28E-09 |
| rs11216178 | 116766390 | T | G | 0.169 | 0.077 | 0.013 | 4.65E-09 |
| rs10892044 | 116766899 | C | T | 0.169 | 0.077 | 0.013 | 4.87E-09 |
| rs7943309 | 116773653 | A | G | 0.064 | 0.115 | 0.020 | 1.27E-08 |
| rs4938317 | 116778275 | C | T | 0.176 | 0.072 | 0.013 | 2.73E-08 |
| rs75920871 | 116780095 | T | A | 0.176 | 0.071 | 0.013 | 3.70E-08 |
| rs117865451 | 116780568 | C | T | 0.065 | 0.115 | 0.020 | 1.15E-08 |
| rs1838757 | 116785083 | C | G | 0.808 | -0.072 | 0.013 | 1.04E-08 |
| rs75487836 | 116789102 | C | T | 0.064 | 0.114 | 0.020 | 1.63E-08 |
| rs7934763 | 116792357 | G | T | 0.177 | 0.070 | 0.013 | 5.73E-08 |
| rs56314469 | 116794903 | A | G | 0.168 | 0.077 | 0.013 | 5.81E-09 |
| rs61907591 | 116796855 | A | T | 0.176 | 0.069 | 0.013 | 9.63E-08 |
| rs61907594 | 116801811 | T | C | 0.175 | 0.069 | 0.013 | 1.01E-07 |
| rs79974099 | 116802581 | A | T | 0.176 | 0.069 | 0.013 | 9.59E-08 |
| rs61907596 | 116807226 | T | C | 0.064 | 0.115 | 0.020 | 1.35E-08 |
| rs61624438 | 116809945 | A | C | 0.176 | 0.070 | 0.013 | 8.63E-08 |
| rs61907602 | 116810655 | C | G | 0.064 | 0.114 | 0.020 | 1.67E-08 |
| rs7941912 | 116815762 | A | G | 0.064 | 0.115 | 0.020 | 1.27E-08 |
| rs7924318 | 116816630 | G | C | 0.176 | 0.069 | 0.013 | 9.37E-08 |
| rs111327557 | 116817784 | T | C | 0.191 | 0.071 | 0.013 | 1.57E-08 |
| rs4316540 | 116819241 | T | C | 0.176 | 0.069 | 0.013 | 1.30E-07 |
| rs61907605 | 116820071 | A | T | 0.176 | 0.069 | 0.013 | 9.61E-08 |
| rs1473324 | 116821918 | G | T | 0.176 | 0.069 | 0.013 | 9.29E-08 |

| SNP[a] | Position on chromosome 11 | Reference allele | Other allele | Effect allele frequency | Beta[b] | Standard error | p-value |
|---|---|---|---|---|---|---|---|
| rs1473325 | 116822034 | T | C | 0.176 | 0.069 | 0.013 | 1.01E-07 |
| rs61907608 | 116823055 | T | C | 0.052 | 0.123 | 0.023 | 4.84E-08 |
| rs1473327 | 116823996 | T | A | 0.176 | 0.069 | 0.013 | 9.71E-08 |
| rs7112967 | 116831912 | T | C | 0.176 | 0.069 | 0.013 | 1.01E-07 |
| rs56134734 | 116833055 | A | G | 0.176 | 0.070 | 0.013 | 6.17E-08 |
| rs61905677 | 116833655 | G | T | 0.176 | 0.069 | 0.013 | 8.86E-08 |
| rs61905678 | 116834785 | T | C | 0.174 | 0.068 | 0.013 | 2.30E-07 |
| rs61905679 | 116834787 | T | C | 0.174 | 0.068 | 0.013 | 1.99E-07 |
| rs11826038 | 116835398 | G | A | 0.064 | 0.116 | 0.020 | 1.17E-08 |
| rs61905680 | 116836031 | A | G | 0.176 | 0.069 | 0.013 | 1.23E-07 |
| rs61905681 | 116836209 | C | A | 0.171 | 0.069 | 0.013 | 1.46E-07 |
| rs61905682 | 116837823 | C | G | 0.064 | 0.115 | 0.020 | 1.49E-08 |
| rs11823013 | 116839535 | C | T | 0.176 | 0.069 | 0.013 | 9.13E-08 |
| rs59294170 | 116841446 | T | A | 0.176 | 0.069 | 0.013 | 9.78E-08 |
| rs17120197 | 116845104 | C | T | 0.219 | 0.055 | 0.012 | 5.02E-06 |
| rs147893781 | 116848716 | T | C | 0.064 | 0.115 | 0.020 | 1.53E-08 |
| rs61905687 | 116862144 | C | T | 0.176 | 0.070 | 0.013 | 8.39E-08 |
| rs7130531 | 116863833 | T | C | 0.176 | 0.069 | 0.013 | 1.09E-07 |
| rs7950213 | 116868654 | A | C | 0.176 | 0.069 | 0.013 | 1.14E-07 |
| rs17120233 | 116871223 | G | A | 0.176 | 0.069 | 0.013 | 1.26E-07 |
| rs75858236 | 116872263 | G | A | 0.176 | 0.069 | 0.013 | 9.96E-08 |
| rs74601470 | 116873382 | T | C | 0.064 | 0.115 | 0.020 | 1.59E-08 |
| rs61905690 | 116876168 | G | C | 0.064 | 0.114 | 0.020 | 1.56E-08 |
| rs77250716 | 116877353 | C | T | 0.176 | 0.069 | 0.013 | 1.27E-07 |
| rs77832134 | 116879636 | G | A | 0.064 | 0.115 | 0.020 | 1.42E-08 |
| rs61905691 | 116880703 | G | A | 0.176 | 0.069 | 0.013 | 1.17E-07 |
| rs7125788 | 116883396 | C | T | 0.176 | 0.069 | 0.013 | 1.17E-07 |
| rs61905704 | 116883665 | C | T | 0.064 | 0.114 | 0.020 | 1.57E-08 |
| rs61905705 | 116883776 | T | A | 0.170 | 0.067 | 0.013 | 3.96E-07 |
| rs61905706 | 116883777 | T | G | 0.170 | 0.067 | 0.013 | 4.76E-07 |
| rs61905708 | 116886159 | T | G | 0.064 | 0.115 | 0.020 | 1.19E-08 |
| rs185955154 | 116888448 | A | G | 0.049 | 0.126 | 0.024 | 1.11E-07 |
| rs4936359 | 116893724 | A | T | 0.178 | 0.069 | 0.013 | 9.75E-08 |
| rs4936360 | 116893769 | C | G | 0.178 | 0.069 | 0.013 | 9.11E-08 |
| rs61905709 | 116897834 | T | C | 0.178 | 0.068 | 0.013 | 1.32E-07 |
| rs4938325 | 116898479 | A | G | 0.178 | 0.068 | 0.013 | 1.17E-07 |
| rs7112191 | 116902056 | A | G | 0.178 | 0.068 | 0.013 | 1.79E-07 |
| rs61905711 | 116906431 | T | C | 0.064 | 0.115 | 0.020 | 1.41E-08 |
| rs61905712 | 116909293 | A | G | 0.178 | 0.068 | 0.013 | 1.19E-07 |
| rs61905713 | 116909429 | T | C | 0.178 | 0.068 | 0.013 | 1.19E-07 |
| rs59266239 | 116911077 | T | C | 0.193 | 0.069 | 0.013 | 3.94E-08 |
| rs61905715 | 116913051 | T | C | 0.064 | 0.115 | 0.020 | 1.53E-08 |
| rs17120241 | 116915020 | C | T | 0.178 | 0.068 | 0.013 | 1.17E-07 |
| rs17496758 | 116916175 | A | G | 0.064 | 0.115 | 0.020 | 1.55E-08 |
| rs61905716 | 116919067 | T | C | 0.177 | 0.067 | 0.013 | 2.11E-07 |
| rs61905717 | 116919208 | A | G | 0.211 | 0.069 | 0.012 | 1.32E-08 |
| rs112151056 | 116919368 | A | G | 0.179 | 0.068 | 0.013 | 1.25E-07 |
| rs17120244 | 116919511 | T | C | 0.212 | 0.069 | 0.012 | 1.13E-08 |
| rs61905718 | 116919677 | T | C | 0.190 | 0.067 | 0.013 | 1.18E-07 |
| rs11826651 | 116920693 | A | T | 0.178 | 0.068 | 0.013 | 1.26E-07 |
| rs61905719 | 116922714 | T | C | 0.178 | 0.068 | 0.013 | 1.14E-07 |
| rs17120251 | 116923488 | C | T | 0.193 | 0.069 | 0.013 | 3.80E-08 |
| rs11826742 | 116924366 | G | C | 0.193 | 0.069 | 0.013 | 3.50E-08 |
| rs723953 | 116924978 | C | A | 0.194 | 0.068 | 0.013 | 4.70E-08 |
| rs17581874 | 116929384 | C | A | 0.064 | 0.115 | 0.020 | 1.40E-08 |
| rs4938333 | 116930888 | G | A | 0.178 | 0.068 | 0.013 | 1.18E-07 |
| rs61903394 | 116931898 | A | G | 0.176 | 0.070 | 0.013 | 8.36E-08 |
| rs61903395 | 116931907 | G | A | 0.191 | 0.070 | 0.013 | 2.75E-08 |

| SNP[a] | Position on chromosome 11 | Reference allele | Other allele | Effect allele frequency | Beta[b] | Standard error | p-value |
|---|---|---|---|---|---|---|---|
| rs112895963 | 116937159 | T | A | 0.177 | 0.068 | 0.013 | 1.46E-07 |
| rs61903398 | 116941613 | T | C | 0.178 | 0.069 | 0.013 | 1.13E-07 |
| rs17120280 | 116942132 | A | G | 0.178 | 0.069 | 0.013 | 8.81E-08 |
| rs61903399 | 116942475 | T | A | 0.178 | 0.069 | 0.013 | 1.10E-07 |
| rs59511712 | 116945762 | T | C | 0.196 | 0.069 | 0.012 | 3.60E-08 |
| rs61903414 | 116946788 | T | C | 0.064 | 0.114 | 0.020 | 1.55E-08 |
| rs4938336 | 116948442 | G | A | 0.178 | 0.068 | 0.013 | 1.16E-07 |
| rs4936362 | 116952200 | T | C | 0.177 | 0.072 | 0.013 | 3.33E-08 |
| rs61903418 | 116956665 | A | G | 0.178 | 0.069 | 0.013 | 9.93E-08 |
| rs61903419 | 116957219 | A | G | 0.211 | 0.069 | 0.012 | 1.12E-08 |
| rs7930409 | 116958943 | G | A | 0.211 | 0.070 | 0.012 | 1.09E-08 |
| rs7930783 | 116959039 | A | G | 0.211 | 0.069 | 0.012 | 1.13E-08 |
| rs111901638 | 116959848 | A | G | 0.178 | 0.069 | 0.013 | 8.64E-08 |
| rs11823889 | 116960403 | A | G | 0.212 | 0.070 | 0.012 | 9.27E-09 |
| rs11823918 | 116960497 | C | G | 0.178 | 0.069 | 0.013 | 9.29E-08 |
| rs56016514 | 116961588 | A | G | 0.212 | 0.069 | 0.012 | 1.04E-08 |
| rs61903421 | 116961802 | T | A | 0.178 | 0.069 | 0.013 | 8.78E-08 |
| rs57582454 | 116961892 | A | G | 0.212 | 0.069 | 0.012 | 1.06E-08 |
| rs61903423 | 116962078 | A | G | 0.211 | 0.070 | 0.012 | 9.82E-09 |
| rs7932655 | 116962563 | A | T | 0.212 | 0.069 | 0.012 | 1.31E-08 |
| rs7946219 | 116962759 | T | C | 0.208 | 0.067 | 0.012 | 4.85E-08 |
| rs7946729 | 116963180 | T | C | 0.212 | 0.069 | 0.012 | 1.07E-08 |
| rs7946869 | 116963312 | T | C | 0.196 | 0.069 | 0.012 | 3.44E-08 |
| rs7950381 | 116963897 | A | C | 0.178 | 0.069 | 0.013 | 8.23E-08 |
| rs7950501 | 116963965 | T | C | 0.178 | 0.069 | 0.013 | 9.48E-08 |
| rs59097294 | 116964437 | C | T | 0.211 | 0.070 | 0.012 | 8.73E-09 |
| rs56227539 | 116964469 | C | T | 0.211 | 0.070 | 0.012 | 8.64E-09 |
| rs61903426 | 116968268 | T | C | 0.211 | 0.060 | 0.012 | 7.33E-07 |
| rs7128071 | 116968798 | A | G | 0.210 | 0.060 | 0.012 | 6.89E-07 |
| rs4936363 | 116972935 | T | C | 0.211 | 0.059 | 0.012 | 8.90E-07 |
| rs11216275 | 116975704 | G | T | 0.274 | 0.051 | 0.011 | 4.10E-06 |
| rs4938339 | 116980594 | T | C | 0.211 | 0.060 | 0.012 | 6.85E-07 |
| rs4938340 | 116980622 | A | G | 0.211 | 0.060 | 0.012 | 8.28E-07 |
| rs6589595 | 116981335 | A | G | 0.212 | 0.061 | 0.012 | 5.18E-07 |
| rs59700451 | 116982928 | A | G | 0.211 | 0.060 | 0.012 | 6.58E-07 |
| rs7113932 | 116985247 | T | G | 0.064 | 0.115 | 0.020 | 1.50E-08 |
| rs7925835 | 116985641 | G | T | 0.064 | 0.114 | 0.020 | 2.32E-08 |
| rs61903429 | 116986684 | A | C | 0.211 | 0.060 | 0.012 | 6.69E-07 |
| rs61903430 | 116987342 | C | G | 0.211 | 0.060 | 0.012 | 6.18E-07 |
| rs75210049 | 116987964 | A | G | 0.063 | 0.114 | 0.020 | 2.89E-08 |
| rs80092204 | 116987988 | C | T | 0.063 | 0.114 | 0.020 | 2.89E-08 |
| rs77511949 | 116988040 | T | A | 0.063 | 0.114 | 0.020 | 2.89E-08 |
| rs75904569 | 116988313 | G | A | 0.208 | 0.062 | 0.012 | 3.05E-07 |
| rs61903459 | 116989185 | G | A | 0.065 | 0.115 | 0.020 | 1.30E-08 |
| rs61905460 | 116989524 | T | C | 0.064 | 0.115 | 0.020 | 1.21E-08 |
| rs7938470 | 116990934 | A | G | 0.064 | 0.116 | 0.020 | 1.17E-08 |
| rs61905462 | 116994106 | A | C | 0.064 | 0.115 | 0.020 | 1.40E-08 |
| rs61905463 | 116997964 | A | G | 0.064 | 0.115 | 0.020 | 1.35E-08 |
| rs61905464 | 117001384 | A | G | 0.065 | 0.115 | 0.020 | 1.32E-08 |
| rs147062051 | 117014926 | A | G | 0.064 | 0.115 | 0.020 | 1.44E-08 |
| rs189193731 | 117015029 | A | C | 0.050 | 0.121 | 0.023 | 1.44E-07 |
| rs181954995 | 117015290 | A | G | 0.064 | 0.113 | 0.020 | 2.21E-08 |
| rs61905472 | 117017706 | T | C | 0.065 | 0.115 | 0.020 | 1.30E-08 |
| rs61905473 | 117018031 | T | G | 0.064 | 0.116 | 0.020 | 1.09E-08 |
| rs61905474 | 117019530 | A | G | 0.065 | 0.115 | 0.020 | 1.22E-08 |
| rs61905475 | 117024435 | G | A | 0.065 | 0.115 | 0.020 | 1.29E-08 |
| rs61905476 | 117024537 | G | A | 0.064 | 0.116 | 0.020 | 1.20E-08 |
| rs61905477 | 117026055 | G | A | 0.065 | 0.115 | 0.020 | 1.29E-08 |

| SNP[a] | Position on chromosome 11 | Reference allele | Other allele | Effect allele frequency | Beta[b] | Standard error | p-value |
|---|---|---|---|---|---|---|---|
| rs12271743 | 117026154 | C | T | 0.065 | 0.113 | 0.020 | 1.96E-08 |
| rs12291885 | 117027942 | C | T | 0.064 | 0.117 | 0.020 | 9.21E-09 |
| rs118155116 | 117035228 | C | T | 0.064 | 0.116 | 0.020 | 1.03E-08 |
| rs61905518 | 117040739 | G | A | 0.065 | 0.115 | 0.020 | 1.28E-08 |
| rs7112577 | 117044603 | G | C | 0.065 | 0.115 | 0.020 | 1.14E-08 |
| rs61905519 | 117045555 | C | T | 0.065 | 0.115 | 0.020 | 1.28E-08 |
| rs77086432 | 117046136 | C | T | 0.065 | 0.115 | 0.020 | 1.29E-08 |
| rs61905521 | 117049020 | A | G | 0.064 | 0.117 | 0.020 | 1.04E-08 |
| rs61905522 | 117049025 | A | G | 0.064 | 0.117 | 0.020 | 1.03E-08 |
| rs61905525 | 117058854 | A | G | 0.064 | 0.115 | 0.020 | 1.23E-08 |
| rs143668212 | 117061197 | T | C | 0.064 | 0.115 | 0.020 | 1.24E-08 |
| rs540723648 | 117061896 | C | G | 0.050 | 0.115 | 0.023 | 5.28E-07 |
| rs61905526 | 117067465 | T | C | 0.064 | 0.116 | 0.020 | 9.51E-09 |
| rs61905527 | 117069966 | T | C | 0.064 | 0.115 | 0.020 | 1.25E-08 |
| rs112333700 | 117071505 | T | C | 0.065 | 0.117 | 0.020 | 7.08E-09 |
| rs117279127 | 117072074 | T | C | 0.065 | 0.116 | 0.020 | 9.31E-09 |
| rs61905530 | 117077716 | A | G | 0.063 | 0.115 | 0.021 | 2.70E-08 |
| rs61906485 | 117082691 | T | G | 0.065 | 0.116 | 0.020 | 1.02E-08 |
| rs61906486 | 117082865 | A | C | 0.065 | 0.115 | 0.020 | 1.19E-08 |
| rs76830818 | 117083831 | T | A | 0.065 | 0.115 | 0.020 | 1.19E-08 |
| rs76604009 | 117085261 | T | C | 0.065 | 0.116 | 0.020 | 8.95E-09 |
| rs61906487 | 117088821 | T | C | 0.064 | 0.117 | 0.020 | 8.54E-09 |
| rs61906488 | 117089505 | C | T | 0.065 | 0.114 | 0.020 | 1.58E-08 |
| rs61906489 | 117091153 | G | C | 0.065 | 0.115 | 0.020 | 1.39E-08 |
| rs148537493 | 117093511 | T | C | 0.052 | 0.117 | 0.022 | 1.86E-07 |
| rs59781045 | 117095283 | T | C | 0.082 | 0.098 | 0.018 | 6.67E-08 |

[a]SNPs with association P values < $7 \times 10^{-6}$ (Bonferroni correction with number of SNPs in the locus) in meta-analysis of five Finnish population cohorts within 1-Mb of the *APOA1* gene are shown; rs12225230 highlighted in grey. [b]Effect sizes (betas) are given as standard deviation (SD) units in inverse-rank normal transformed covariate-adjusted metabolite residuals.

**Supplemental Table 7. Association of rs12225230 with conventional vascular risk factors.**

| Study | Phenotype | Sample size | Effect (beta and se) for minor allele of rs12225230 | *p* value |
|---|---|---|---|---|
| UK Biobank[31] | Alcohol intake frequency | 336,965 | -0.0072 (0.0047)[b] | 0.124 |
| | Body mass index | 336,107 | -0.0022 (0.0032)[b] | 0.494 |
| | Smoking status: current | 336,024 | 0.0015 (0.00097)[c] | 0.111 |
| | Systolic blood pressure | 317,754 | 0.0003 (0.0032)[b] | 0.938 |
| DIAGRAM[32] | Type 2 diabetes | 159,208 | 0.011 (0.016)[c] | 0.470 |

[a]The phenotype associations were screened from publicly available GWAS data using PhenoScanner v2[15]. [b]Beta for inverse normally rank transformed value. [c]Logarithm of odds ratio.

**Supplemental Table 8. Observational associations of circulating apoA-I, apoB, HDL cholesterol, and LDL cholesterol concentrations with coronary artery disease.**

| Model[a] | Metabolic measure | HR (95% CI)[b] | *p* value |
|---|---|---|---|
| Minimally adjusted | apoA-I (higher) | 0.82 (0.76-0.88) | 2.0E-07 |
| | apoB (lower) | 0.81 (0.75-0.87) | 7.0E-09 |
| | HDL-C (higher) | 0.75 (0.69-0.81) | 1.6E-12 |
| | LDL-C (lower) | 0.87 (0.81-0.94) | 3.0E-04 |
| Fully adjusted | apoA-I (higher) | 0.86 (0.79-0.93) | 2.0E-04 |
| | apoB (lower) | 0.84 (0.79-0.91) | 7.0E-09 |
| | HDL-C (higher) | 0.79 (0.72-0.86) | 4.4E-06 |
| | LDL-C (lower) | 0.88 (0.82-0.95) | 6.3E-04 |

[a]The observational associations were estimated in the FINRISK97 cohort (*n*= 7,133, number of CAD events= 743) under fully (age, sex, body-mass index, systolic blood pressure, type 2 diabetes, smoking and alcohol consumption as covariates) and minimally (age and sex as covariates) adjusted models. [b]The effects are shown per 1-SD higher (apoA-I, HDL-C) or lower (apoB, LDL-C) concentration.

**Supplemental Table 9. Effect of apoA-I infusion therapies on risk of major adverse cardiovascular events.**

| Trial | Drug | Dose[a] | *n* (%) of MACE cases treatment / placebo | Total *n* treatment / placebo | RR of apo-AI vs placebo (95% CI) |
|---|---|---|---|---|---|
| CHI-SQUARE[28] | CER-001 | 3 mg/kg (~0.27 g) | 16 (13.3) / 10 (8.3) | 120 / 120 | 1.60 (0.76-3.38) |
| | | 6 mg/kg (~0.54 g) | 17 (13.7) / 10 (8.3) | 124 / 120 | 1.65 (0.79-3.45) |
| | | 12 mg/kg (~1.07 g) | 12 (9.8) / 10 (8.3) | 122 / 120 | 1.18 (0.53-2.63) |
| AEGIS-I[29] | CSL112 | 2 g | 27 (6.4) / 23 (5.5) | 419 / 418 | 1.17 (0.68-2.01) |
| | | 6 g | 24 (5.7) / 23 (5.5) | 421 / 418 | 1.04 (0.59-1.81) |

[a]For CHI-SQUARE, the approximate doses in grams (in parentheses) were estimated based on mean weights reported in the study.

**Supplemental Table 10. Association of rs12225230 with cardiovascular disease in large GWAS consortia.**

| Study | Outcome phenotype | Sample size | Effect (beta and se) for minor allele of rs12225230 | *p* value |
|---|---|---|---|---|
| Nikpay *et al.* 2015[33] / CARDIoGRAMplusC4D Consortium[a] | Coronary artery disease | 184,305 | 0.0156 (0.0118) | 0.186 |
| | Myocardial infarction | 171,875 | 0.0178 (0.0130) | 0.171 |
| Nelson *et al.* 2017[23] / UK Biobank, CARDIoGRAMplusC4D, Myocardial Infarction Genetics and CARDIoGRAM[b] | Coronary artery disease | 332,495 | 0.0117 (0.0104) | 0.261 |
| van der Harst *et al.* 2018[22] / UK Biobank and CardioGRAMplusC4D[c] | Coronary artery disease | 547,261 | 0.0124 (0.0073) | 0.090 |
| Malik *et al.* 2018[25] / MEGASTROKE[d] | Stroke[e] | 521,612 | -0.0146 (0.0106) | 0.169 |

[a]Meta-analysis of GWAS studies of mainly European, South Asian, and East Asian descent; [b]Meta-analysis of UK Biobank SOFT CAD GWAS (interim release) with CARDIoGRAMplusC4D 1000 Genomes-based GWAS and the Myocardial Infarction Genetics and CARDIoGRAM Exome; [c]Meta-analysis of UK Biobank and CardioGRAMplusC4D; [d]Meta-analysis of 29 studies (trans-ethnic); [e]Any stroke (comprising ischemic stroke, intracerebral hemorrhage, and stroke of unknown type).
